# The Sniffbot: A biohybrid robot for active sensing-based odor localization and discrimination

**DOI:** 10.64898/2026.02.17.706375

**Authors:** Neta Shvil, Noa Gozin, Anton Sheinin, Omer Yuval, Yossi Yovel, Ben Meir Maoz, Amir Ayali

## Abstract

The detection, identification and localization of volatile compounds are of critical importance for various applications, ranging from gas leak detection to drug and explosive sensing. Current technologies—such as gas chromatography–mass spectrometry and e-noses—are limited by slow analysis, low mobility, and reduced sensitivity and adaptability, making them unsuitable for real-time odor localization in real-world settings. Here, we present ‘Sniffbot’: an autonomous, mobile biohybrid robotic sensory system that overcomes these challenges by harnessing the extraordinary olfactory capabilities of the desert locust antenna, an advanced olfactory sensor, that generates odorant-specific electrophysiological responses to numerous odorants. Our Sniffbot platform consists of a compact robotic vehicle onto which we have assembled: (i) a sensing module, comprising a locust antenna and a miniaturized electrophysiology system; (ii) a “sniffing” module, which actively samples air in the environment, creating a timed airflow over the antenna, preventing the antenna from becoming habituated to odorant stimuli; and (iii) a decision-making module that analyzes the sensory input in real time to navigate or identify odors. Sniffbot’s movements are controlled by an odorant-search algorithm coupled with the sniffing module. This enables Sniffbot to detect and localize odors independently of wind-induced odorant gradients, and thus to be used in challenging windless environments. The Trident, a novel search algorithm, outperforms several commonly used algorithms in localizing the odorant source. We further demonstrate Sniffbot’s ability to discriminate a target odor among others. Our results demonstrate the potential of augmenting biological sensors with autonomous robotic components for next-generation chemical sensing and environmental monitoring.

## 1. Introduction

The capacity to detect the presence of volatile compounds (odorants), identify them, and localize their sources is critical to many applications, including detection of gas or chemical leaks, drugs, explosives, food spoilage, and more [1–6]. Several technologies are currently available for odorant detection and identification, such as gas chromatography-mass spectrometry (GC-MS) and recently developed electronic sensors (e-noses). Yet these technologies fall short in terms of real-world applicability. In particular, they are not sufficiently mobile or autonomous to localize odors in real time: For example, GC-MS relies on the collection and laboratory-based analysis of air samples, and thus cannot perform real-time, on-site odor tracking, or operate effectively in dynamic, unstructured environment [7,8]. Similarly, e-noses do offer portability and faster analysis but cannot match the sensitivity, selectivity, or contextual adaptability of biological olfactory systems [8–10]. Because of these shortcomings, animals such as sniffer dogs are commonly used in time-sensitive applications requiring odorant detection and localization [11]. Yet, these setups are also limited: dogs are trained to respond to a small set of target odorants. Their training is lengthy and expensive and thus their numbers are limited, and their performance can be influenced by environmental conditions such as humidity and temperature [12]. Here, we present a biohybrid robotic sensing platform that overcomes these challenges by exploiting the natural odorant-sensing capabilities of animals (insects). This robot is composed of a biologically inspired active-sensing, a machine-learning based discrimination system and an ability to search for an odor, all integrated into a compact autonomous mobile platform **(Figure. 1A&B, Supp. Figure. 1)**.

**Figure 1:**
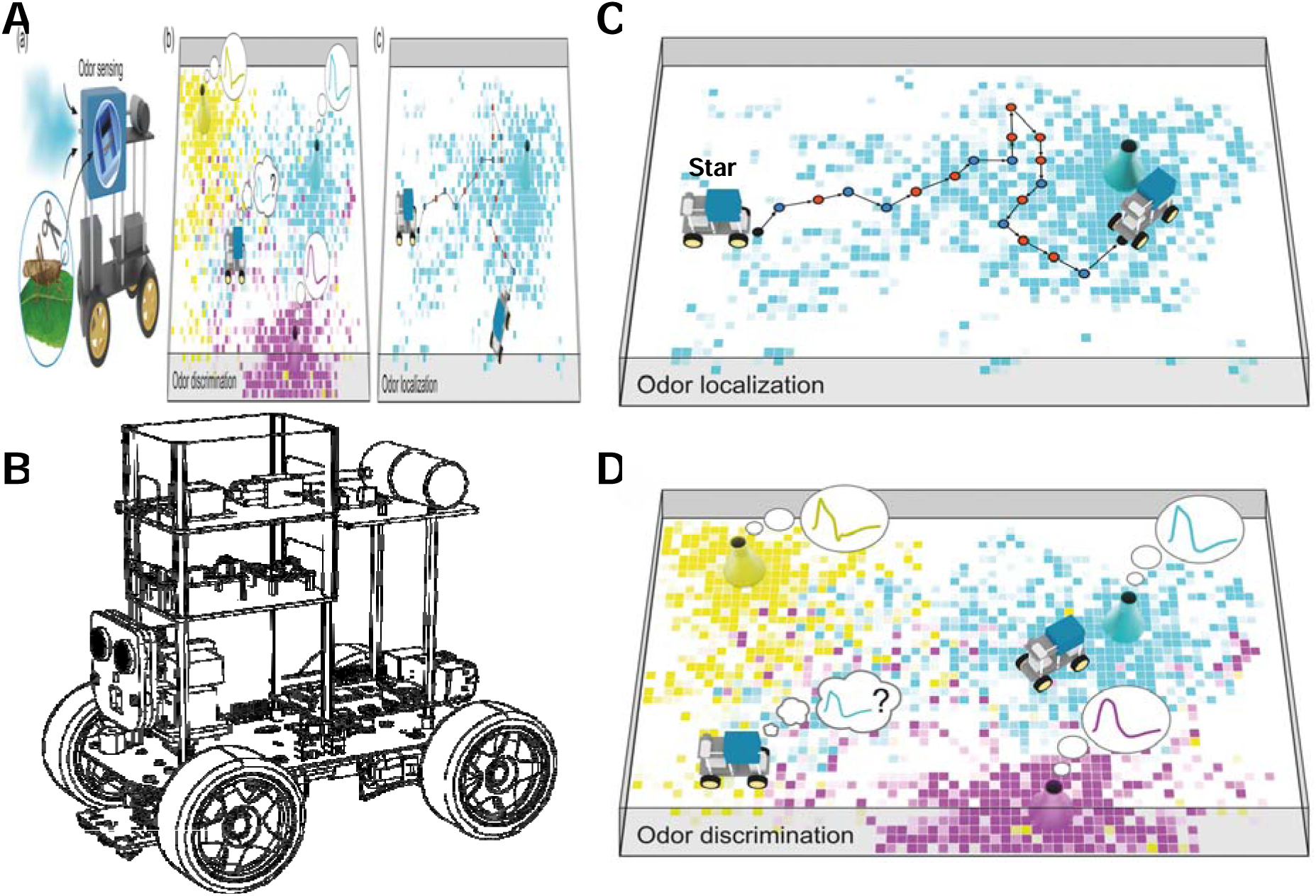
Sniffbot: A Bio-Hybrid Robot for Odor Localization and Discrimination. **(A)** The antenna of the desert locust is removed and placed in the antenna holder within Sniffbot, which employs active sensing to sample the surrounding air for odor cues. **(B)** The assembly of Sniffbot, illustrating its three integrated layers. The sniffing and sensing modules, described in the paper, are shown in color: the sniffing layer is on top, and the sensing layer is directly beneath it. The decision-making layer and the robotic chassis are shown in black and white. **(C)** Sniffbot uses odor cues to navigate toward an odor source. **(D)**Sniffbot uses a machine learning model to discriminate between different odors.

The core sensory component of our system is the odorant sensor, an olfactory organ (antenna) of an insect (**Figure. 1A**). Researchers have long acknowledged insects’ promise in odorant-sensing applications, given their superior olfactory capabilities [13–17]: For example, the female mosquito can sense CO_2_ from a 10m distance [18], the male moth can respond to a single molecule of the female’s sex pheromone and follow its location over distances greater than 80m [16,19], and fruit flies have been reported to identify acetic acid at concentrations lower than a part per million (PPM) [20]. Alongside these capabilities, insects’ sensing organs (mainly antennae) are amenable to integration in biohybrid platforms [14,21,22], given their small size, high sensitivity and low energy consumption [23,24]. Particularly important for our context, insect antennae can generate odorant-specific electrophysiological signals that can be measured with an electroantennogram (EAG) and used to identify and discriminate between individual or mixtures of odorants [9,25–27]. We and others have recently shown that EAG signals generated by the locust antenna—even when detached from the insect [7]—can be used to detect and discern a large variety of odorants, with high sensitivity [9,13,15]. In this study, we developed a miniaturized EAG system that can be mounted onto a small robotic vehicle to measure signals produced by an insect antenna in real time. We further show that this system, in conjunction with an autonomous machine-learning classifier, successfully identifies and discerns among multiple types of odorants.

One of the greatest challenges in odorant-sensing technology is effective localization of an odorant source [28]. For many animals, odor localization is dependent on the presence of wind, which creates odorant plumes that signal the directionality of the odor source (i.e., upwind). In a process called anemotaxis, animals use various search strategies to move towards the odor source according to wind direction [28–33]. Several odorant localization technologies rely on similar principles, that is, measuring chemical gradients resulting from air flow and moving accordingly [34]. Yet, reliance on air movement for odorant localization creates challenges in windless or nearly-still settings, such as indoor rooms, industrial warehouses, or collapsed-building interiors, where environmental airflow is typically under 0.1Lm/s [34–36]. In the absence of wind, odorants primarily spread through diffusion, generating weak, shallow gradients that are difficult to resolve. In these conditions, the diffusion-induced gradient of odorant concentration is typically limited to a small area around the odor source (up to 40 cm) [31,37,38]. Beyond this zone, residual air currents (velocities <0.05 m/s) outpace molecular diffusion, creating randomly distributed “air pockets” or patches [31,37], and yielding no exploitable concentration gradients for plume following. This creates a complex-dynamic landscape in which the presence or concentration of odorant molecules provides little information regarding the location of the odorant source [31,34,39,40]. Researchers have developed several search algorithms to localize odorant sources under such conditions, including the E. coli (random walk), Spiral, and Hex path (zigzag) algorithms [34,39,40]; to our knowledge, none of these have been deployed in biohybrid systems. More advanced strategies include information-theoretic approaches such as infotaxis and Bayesian belief updating [41–43], finite-state controllers with limited memory [44], reinforcement-learning policies for turbulent plumes [45], computational modeling [46], and multi-robot swarm strategies [47,48]. However, many of these assume wind-driven plumes, continuous concentration measurements, or substantial computational resources, making them challenging for resource-constrained biohybrid platforms and for windless indoor environments. Here, we develop a new search algorithm (Trident) that, when used in combination with a biological sensor (an insect’s antenna), outperforms classical algorithms in its capacity to localize odorant sources in windless conditions. All algorithms in this study were tested under windless conditions to evaluate their effectiveness in the most challenging scenario for odor localization. Another challenge introduced by windless environments is specific to biological sensors: Sensory neurons tend to become habituated (i.e., to show a decreasing activation response) to continuous or repeated stimuli (such as an odor in a windless environment), and to respond more robustly to transient stimuli [49–52]. We prevent such habituation by introducing an active air-sampling module that mimics biological sniffing by creating a timed airflow over the antenna (**Fig 1A&B**, **Figure. 2B&C, Supp. Figure. 2A, Movie. 1)**.

**Figure 2:**
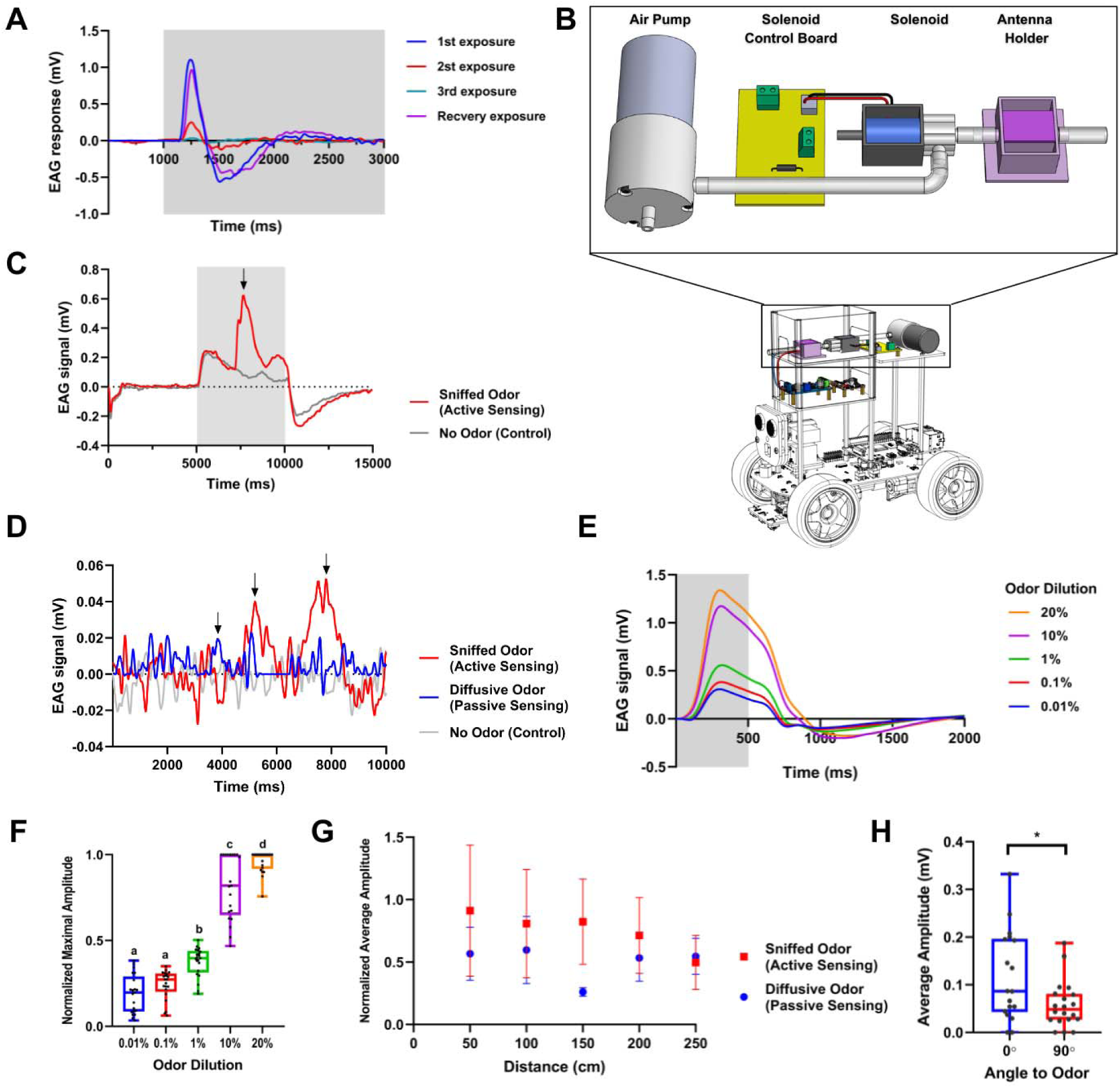
Sniffing Module Benefits, Dose-Dependent Odorant Sensing and Distance Discrimination. **(A)** Antenna habituation experiment: The stimulus period is represented by the gray square. N = 8. The habituation experiments were performed on a detached antenna using the stationary odor delivery system described in Shvil et al. [9] **(B)** Schematic of the sniffer prototype. **(C)** Example of an electroantennogram (EAG) active signal recorded by the sniffer over 5 seconds. The antennal response to an odor-free environment is shown in gray, with the detected peak indicated by a black arrow, and the sniffing period (open solenoid) represented by a gray square. **(D)** Examples of EAG sniffed odor (passive) and diffusive odor (active) signals recorded 100 cm from the odor source. The antennal response to an odor-free environment is displayed in gray, with detected peaks marked by black arrows. **(E)** Averaged EAG signal for different odor concentrations. 500 milliseconds of exposure within a 2000-millisecond recording period. The stimulus period is indicated by the gray square, N=7 antennae and 21 repetitions. **(F)** Maximum point of the normalized EAG signal for different odor concentrations. Letters above the columns denote statistical groupings in two-tailed t-tests. N=7 antennae and 21 repetitions. **(G)** Normalized average peak amplitude of the EAG signal as a function of Sniffbot’s distance from the odor source. Active sniffing (red): significant correlation despite patchy plumes. Passive (blue): flat response due to antennal habituation (Pearson correlation test: r = -0.90, p = 0.03 for sniffed odor and r = -0.2, p = 0.74 for diffusive odor). N=10 antennae; 41 repetitions for passive experiments and N=9 antennae; 25 repetitions for active experiments. Vertical lines represent standard error. **(H)** Average peak amplitude in the sniffer directionality experiment. N=6 antennae, 18 repetitions. Two-tailed t-test: p<0.05.

This work presents a biohybrid olfactory robotic platform that combines active sniffing, a broad-spectrum biological antenna, and rapid temporal dynamics to enable real-time odor identification and localization in windless environment (<0.025 m/s). The resulting “sniffing robot”, Sniffbot, is a compact, autonomous biohybrid mobile sensor with the capacity to detect, identify, and localize a large variety of odorants in real time, in diverse practical settings that have previously been inaccessible to current technologies.

## 2. Materials and Methods

### 2.1 Animals

In all experiments, we used adult desert locusts (*Schistocerca gregaria*) of both sexes. Gregarious phase locusts were raised in a 60-liter metal cage at a density of around 100 insects per cage in our breeding colony at the School of Zoology, Tel Aviv University. The locusts were maintained under controlled conditions with a 12-hour dark/light cycle, temperatures of 35–37°C during the day (with additional heat provided by 25-watt incandescent bulbs) and 30°C at night, and 35–60% humidity. Their diet consisted of wheat grass and dry oats. All experimental procedures were conducted in accordance with accepted regulations and guidelines for research with invertebrates, which do not require formal ethical approval. Experimental procedures were conducted swiftly and gently to minimize handling time and potential distress after which locusts were returned to their rearing cages.

### 2.2 Electroantennogram Recordings

EAG recordings were performed as described previously by Shvil et al. [9]. Briefly, the locust antenna is carefully excised, its distal end trimmed and then placed in an antenna holder so that both cut ends are immersed in an electroconductive gel, while the central portion remains exposed to air (**Figure. 1A, Supp. Figure. 1B**). Two silver electrodes (0.025″ bare, 0.030″ coated, PFA silver (Ag); A-M Systems, Sequim, WA) are used to record the differential signal. The signal is amplified (×100) using a custom-made amplifier (**Supp. Figure. 1C**), filtered (0.5 Hz high-pass/500 Hz low-pass), and finally sampled at approximately 60 Hz. A 4-channel ADC (SparkFun Qwiic 12 Bit ADC; **Supp. Figure. 1D**) converts the analog signal to digital form and transmits it to a Raspberry Pi-4 control board via a customized acquisition code written in Python 3 (Python Software Foundation, Scotts Valley, CA).

### 2.3 Odorants

Lemon oil (NOW Foods, essential oils, lemon) diluted 1:10 (10%) in mineral oil (Sigma-Aldrich) was used in most experiments. For the dose-response experiment, lemon oil was further diluted to 20%, 10%, 1%, 0.1%, and 0.01% to achieve a total volume of 15 mL. For the mapping, distance (active vs. passive sensing), and localization experiments, 2 mL of 10% lemon oil was placed in a candle burner. In the discrimination experiments, 200 µL of either Benzaldehyde or β-Citronellol (Sigma-Aldrich), both diluted to 93 mM, was applied to a round filter paper (Whatman #41, WHATMAN LIMITED, England), which was then presented in front of the sniffer. The directionality experiment was conducted in the same manner but using 10% lemon oil.

### 2.4 Habituation Experiments

The habituation experiments were performed using 93 mM Benzaldehyde in the stationary odor delivery system as described in Shvil et al [9]. The antenna was exposed to the odor for 100 seconds, three times consecutively with a 1-second interval between exposures. It was then exposed once more after a 30-second recovery period.

### 2.5 Data Analysis and Peak Detection

Recording data were sampled using an ADS1015 analog-to-digital converter (Texas Instruments) with an Adafruit module at 60 Hz. After acquisition, the data were processed as follows: (1) the data were interpolated to 1000 Hz using cubic spline interpolation, (2) the signal was offset by subtracting its median value, (3) a digital low-pass filter with a cutoff frequency of 5Hz was applied using the SciPy module command: *signal.butter(4, 5, btype=’lowpass’, fs=1000, output=’sos’)* and (4) the processed signal was then analyzed for odor peaks using the *find_peaks* function from the SciPy *Signal* module with the following four parameters:

- Prominence: the minimum relative amplitude (above the baseline) required for a peak. Baseline is determined automatically by the module.
- Width: the minimum width of a peak at its relative height. Setting a minimum width helps to distinguish true odor-triggered signals, which are typically wider, from noise.
- `o` Both baseline and width are illustrated by the orange and green horizontal lines in **Supp. Figure. 3A**&**B**
- Distance: The minimum time interval between two consecutive peaks, indicated by the purple horizontal line in **Supp. Figure. 3A**&**B**. This parameter is fixed at 1000.
- Relative Height: This parameter is set to a constant value (0.5).

**Figure 3:**
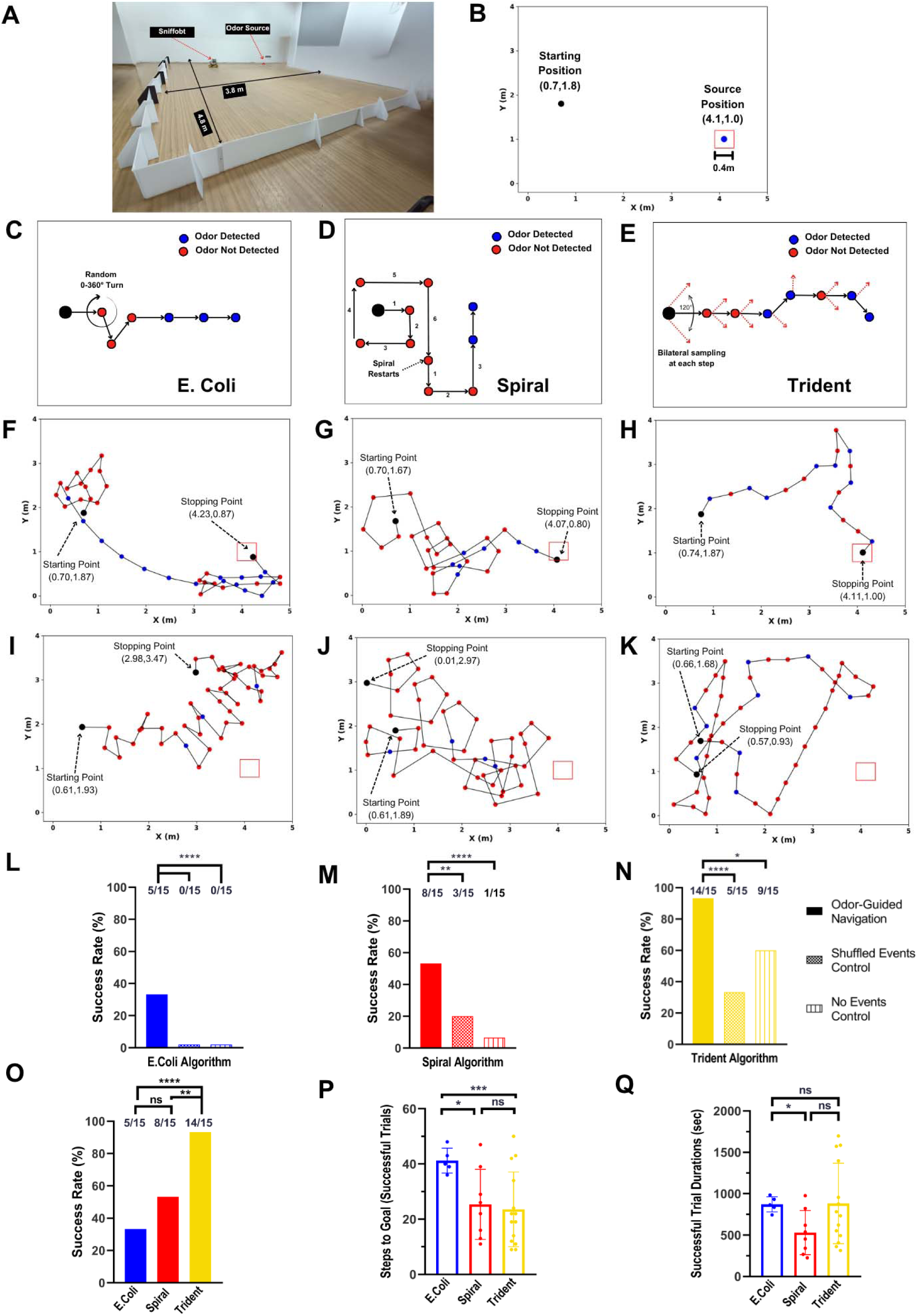
Sniffbot’s Odor Localization Capabilities. **(A)** Assembled photograph of the experimental site. **(B)** Schematic representation of the experimental site**. (C, D, & E)** Schematic trajectories of Sniffbot using the E. coli, Spiral, and Trident algorithms, respectively. **(F, G, & H)** Trajectories from successful experiments using the E. coli, Spiral, and Trident algorithms, respectively. **(I, J, & K)** Trajectories from unsuccessful experiments using the E. coli, Spiral, and Trident algorithms, respectively. The odor source is indicated by a red square in panels (C–K). **(L, M, & N)** Success rates of odor-guided navigation versus two types of odorless controls for the E. coli, Spiral, and Trident algorithms, respectively, when experiments are limited to 50 steps (N = 15). P-values in two-tailed binomial test for E.coli p<0.0001, for Spiral p<0.0001 (no-event control) and p<0.01 (shuffled event control), for Trident p<0.05 (no-event control) and p<0.0001 (Shuffled event control)) **(O)** Overall success rate of odor-guided navigation across all three algorithms. Two-tailed binomial test: p < 0.0001 between Trident and E. coli, p<0.01 between Trident and Spiral. **(P & Q)** Number of steps (P) and time (Q) required for Sniffbot to reach the odor source in successful trials across all three algorithms. Two-tailed t-test between Spiral and E.coli: p < 0.05. Welch’s test between E. coli and Trident: p < 0.0005.

The prominence and width parameters were adjusted anew for every antenna at the beginning of each experiment based on the control (no odor) signals recorded. This was necessary because different antennae may vary in sensitivity thresholds. Given the bipolar nature of the signal, only its positive component was analyzed for peak detection. This approach set the automatically selected baseline to zero, thereby preventing false positives in peak identification (**Supp. Figure. 3**).

### 2.6 Localization: Experimental Design

Localization experiments were conducted in a closed room (doors and windows kept shut) with air conditioning off to simulate a windless indoor environment. Air velocity in the room was consistently below 0.025 m/s on average, in agreement with commonly accepted definitions of windless conditions (measured at ∼23.5°C using SwemaAir40, SWEMA, Sweden). These measurements confirm that residual air motion was well below thresholds typically required for plume formation. Ambient temperature was maintained between 21 °C and 28 °C. Between trials, the room was cross ventilated for at least 30 minutes to remove residual odors. Experiments took place in a 3.8 m × 4.8 m rectangular arena bounded by walls at Sniffbot’s height (**Figure. 3A&B**). An obstacle-avoidance routine ran on Sniffbot, in which ultrasonic sensors measured the distance to the border of the arena before each step and turned away if a wall was detected close by.

At the start of each experiment, a fresh antenna was inserted into the antenna holder and its functionality was verified by presenting an odor vial at the sniffer end during a preliminary sniffing test. Following a successful positive control, three samples of fresh ambient air were taken to determine the minimum threshold and time-frame settings at which no (false-positive) peaks were detected. Next, 2mL of odor was applied to the burner and a candle was lit underneath. The odor was allowed to disperse for ten minutes, after which the candle was extinguished. Sniffbot was then positioned at the starting point, and a remote-controlled code (connected *via* Wi-Fi) was initiated. Each sampling round lasted 15 seconds, with the first two seconds excluded from peak analysis to avoid false-positive detections caused by the antenna’s response to the initial wind burst **(Supp. Figure. 6A).** The experiment either concluded automatically, when Sniffbot reached the predetermined step quota, or manually by the experimenter upon visual confirmation that it had entered the odor source 40x40 cm^2^ square. At the end of each experiment, a second EAG sensing positive control was performed to ensure the antenna is still responsive and validates the results. Only experiments in which both the initial and final positive controls were successful were included in the analysis. All experiments were recorded using a GoPro Hero6 camera for subsequent trajectory analysis.

### 2.7 Robot Trajectory Analysis and Mapping Data

For each experiment, the robot’s trajectory was analyzed and visualized using the following workflow. First, the experimental video was imported into an analysis script written in Python 3 (Python Software Foundation, Scotts Valley, CA) and cropped to match the arena boundaries **(Supp. Figure. 4A).** Sniffbot’s initial position was manually determined on a high-contrast frame using a white sticker on top of the robot as a reference **(Supp. Figure. 4B).** The script then recorded the robot’s coordinates every 12 frames, which were subsequently plotted as a scatter plot **(Supp. Figure. 4C).** These data points were clustered according to the number of steps in the experiment, and the average coordinates of each cluster were color-coded based on whether odor was detected at that location **(Supp. Figs. 4D**&**E)**. Finally, each trajectory was manually validated, with minor adjustments made if necessary.

**Figure 4:**
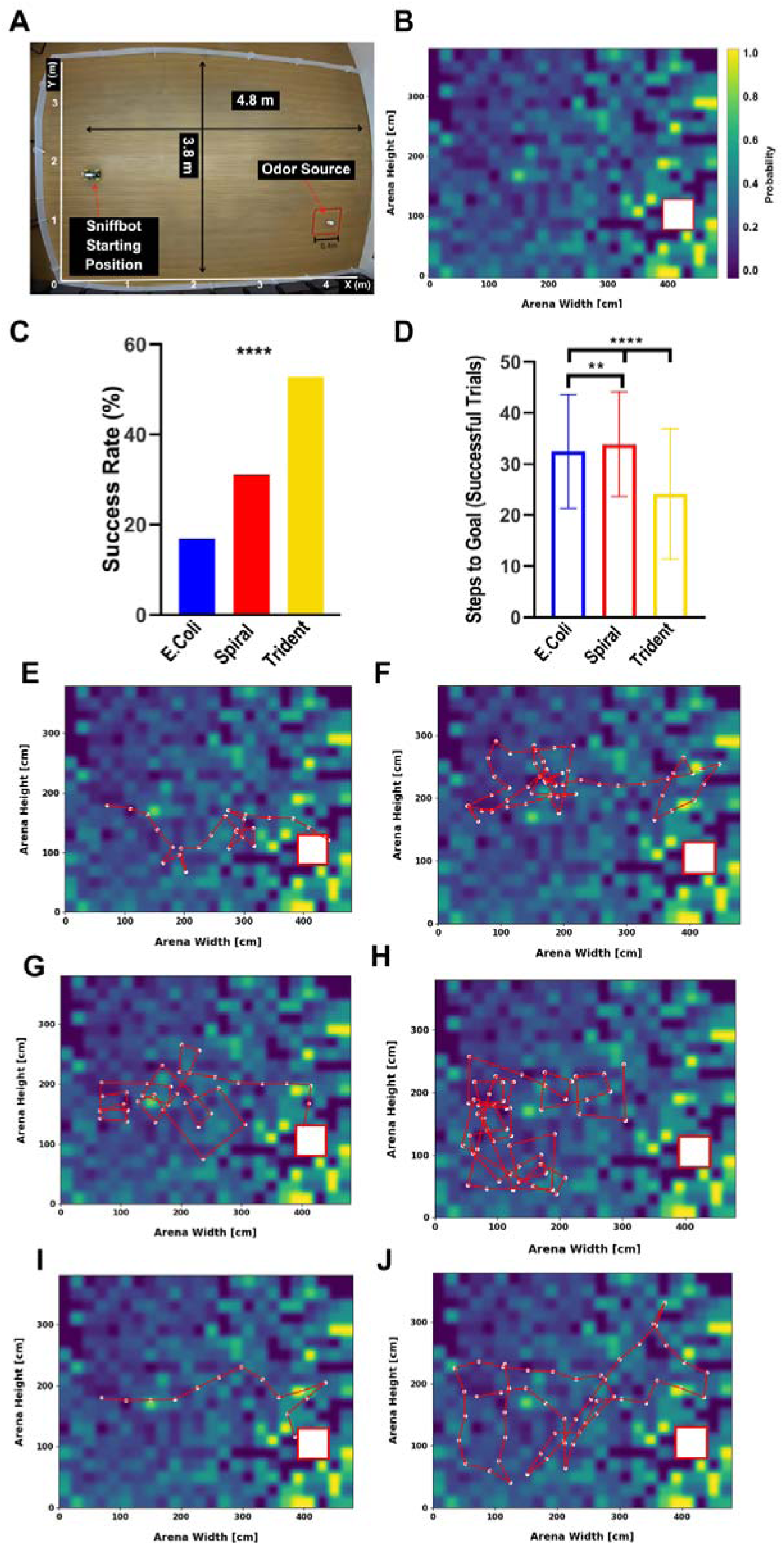
Odor Dispersal in a Windless Environment and Odor-Localization Simulations. **(A)** Photograph of the experimental site with dimensions indicated. **(B)** Heatmap showing the probability of encountering odor based on Sniffbot measurements (N = 63 antennae, 3957 data points). The heatmap is presented with Gaussian smoothing (sigma factor = 4). The odor source is marked by a red square. **(C)** Success rates from simulated experiments for odor-guided navigation across all three algorithms. Two-tailed binomial test: p<0.0001, n=1000. **(D)** Number of steps required for Sniffbot to reach the odor source in successful simulated trials for each algorithm. Two-tailed t-test: p<0.01 and p<0.0001, n=1000. **(E, G & I)** Simulated trajectories of successful experiments using the E. coli, Spiral, and Trident algorithms, respectively. **(F, H, & J)** Simulated trajectories of unsuccessful experiments using the E. coli, Spiral, and Trident algorithms, respectively. All simulations were conducted on the smoothed odor dispersal map (panel B), which was derived from actual odor dispersal measurements made by Sniffbot.

For odor mapping, all coordinate points and their corresponding odor detection labels were used to calculate the probability of odor encounter at each location, normalized by the number of measurements taken at that point. The map was smoothed using Gaussian smoothing (sigma factor = 4) (**Figure. 4B**).

### 2.8 Simulations

Simulations were implemented in Python 3.11 to model the trajectory of the robot in odor-assisted navigation. The odor map was based on real-world experimental data, providing realistic spatial gradients for navigation (**Figure. 4B**). At each step, the code generated a random number and compared it to the probability of encountering odor at that location, as derived from the field experiments. This served as the threshold for determining odor detection. Each algorithm was simulated 1,000 times, with a step limit of 50 (matching the field experiments). The program automatically collected performance metrics, such as success rate and number of steps to goal (in successful trials), and generated summary graphs to visualize statistical outcomes across trials. Core libraries used include NumPy, Pandas, and Matplotlib.

### 2.9 Odor Discrimination: Experimental Design

All the discrimination experiments were conducted using an improved sensing apparatus (see **Supplementary Information**), according to the following procedure: Three filter papers were positioned at Sniffbot’s height—two containing odorants and one serving as a blank control—placed approximately 40 cm from the robot and evenly spaced from one another. The positions of the odorants alternated between experiments. After securing a fresh locust antenna in the antenna holder on Sniffbot, a positive control was performed using lemon oil. Following a five-minute acclimation period to allow the antenna to adjust to the wind conditions, the experiment commenced according to the procedure outlined in **Supp. Figure. 6I**. Each antenna was tested for all three odors of interest in three separate trials. At the end of each experiment, a second positive control was performed to confirm the viability of the antenna. As with the localization experiments, all discrimination tests were conducted in a closed room without air conditioning, with ambient temperatures maintained between 21–28°C. All experiments were recorded on video using a GoPro Hero6.

### 2.10 Statistical Analysis

To evaluate the correlation between the distance from the odor source and the average peak amplitude in both active and passive sensing, we employed the Pearson correlation test. For the dose-response and directionality experiments, a two-tailed t-test was used to assess significance. To compare the success rates of the three navigation algorithms, in both the simulations and field experiments, we applied a two-tailed binomial test. Here, the expected chance was defined either as the success rate in the control experiment (when comparing an algorithm to its controls) or as the success rate in the less successful experiment (when comparing two different algorithms).

The durations of successful trials and the number of steps required to reach the goal were compared using a two-tailed t-test, except for the comparison between the E. coli and Trident algorithms, for which Welch’s test was used due to unequal variances. The discrimination success rates were analyzed using a two-tailed binomial test, with the expected chance defined as the random probability among three choices. The comparison of the predicted probabilities was performed using a two-tailed t-test. Finally, to assess the significance of the classifier’s performance, we used a two-tailed binomial test, where the expected chance was defined by the number of classes and N represented the number of antennae used in each experiment.

## 3. Results

### 3.1 Design and Construction of Sniffbot

The autonomous Sniffbot comprises three main modules—(i) sensing; (ii) sniffing; and (iii) decision-making **(Figure. 1A&B, Supp. Figure. 1)**—assembled onto a commercially available robotic vehicle (21cmX15cmX11cm), equipped with a small single-board Raspberry Pi computer (Freenove 4WD Smart Car Kit for Raspberry Pi; Freenove Creative Technology Co., Ltd & Raspberry Pi) **(Supp. Figure. 1)**. The mobile robotic platform is also equipped with ultrasonic sensors that enabled it to avoid large physical obstacles in its path.

The *sensing* module includes a custom-made antenna holder into which a desert locust antenna and two silver electrodes are inserted (**Figure. 1A&B, Supp. Figure. 1B**). The special design is aimed at retaining the antenna’s viability for several hours and enhancing the signal-to-noise ratio (SNR) of the electrophysiological recordings, as demonstrated and well-established in our previous study^9^ (**Supp. Figure. 1**). The electrodes are connected to a custom-built miniaturized amplifier as part of a miniaturized EAG recording system for signal acquisition, also comprising a digital-to-analog converter and a Faraday cage (**Supp. Figure. 1C, D&E**).

The *sniffing* module is designed to overcome the tendency of many biological sensors to become habituated to prolonged and static stimuli [49–53]. Indeed, in early experiments that we have conducted (see **Materials and Methods**) the locust antennae exhibited habituation (a decrease in activation) during continuous repeated exposures to an odor (1 sec intervals, **Figure. 2A**). However, recovery time was short (∼30 s, **Figure. 2A**). To meet this challenge, we created an active air sampling module that mimics biological sniffing. A solenoid, connected to a small pump, directs airflow through the antenna holder, which was designed to be airtight with a two-lid system (**Figure. 2A, Supp. Figure. 2A**). This setup allows precise, timed airflow over the antenna via the Raspberry Pi (**Fig 1A&B**, **Figure. 2B&C, Supp. Figure. 2A, Movie. 1)**. The sniffing action concentrates odor molecules so that more molecules reach the antenna simultaneously, effectively amplifying the response. Our EAG recordings confirm this effect, showing that actively sniffed odor stimuli produce peaks of higher amplitude compared to passively diffusive odor stimuli (**Figure. 2D**). Details of the peak detection procedure are provided in **Materials and Methods** and in **Supp. Figure. 3**.

The decision-making module, implemented on an on-board computer (Raspberry Pi), analyzes sensory input in real time and generates motor commands accordingly. During navigation tasks, it relies on pre-integrated bio-inspired algorithms, while in discrimination tasks, it utilizes a machine learning classifier to identify odor types. The resulting decisions are translated into movement instructions for the robot.

### 3.2 Dose-Dependent Odorant Sensing and Distance Discrimination

We first conducted a set of experiments to confirm the previously reported dose-dependent odorant sensing behavior of the locust antenna [9]. The EAG signals followed a clear dose-response curve (20%-0.01% odor dilutions; **Figure. 2E, F**).

Next, we assessed the capacity of the sensing module to encode odorants’ distance (i.e., source distance). To this end, we conducted a set of experiments exposing it to an odor at increasing distances from the source (50-250cm; **Figure. 2G, Supp. Figure. 2B-D**). We averaged the amplitudes of all peaks detected in each recording and then assessed the correlation between these averages and the distance from the source; this was done with and without the use of the sniffer module (see **Supplementary Information**). When the sniffer module was used for sampling, we observed a significant negative correlation between the signals’ average amplitude and the distance from the source (Pearson: r = - 0.90, p = 0.03). No such correlation was observed when the EAG measurements were conducted without the sniffer (r = -0.2, p = 0.74) (see also **Supp. Figure. 2B**). Analysis of individual antennae further confirms that this effect is not attributable to averaging (**Supp. Figure. 2D)**. In addition to the distance sensitivity, Sniffbot provides directional (angular) information about the odor source. As shown in **Figure. 2H, Supp. Figure. 2E**, significantly higher average peak signal amplitudes were recorded when the sniffing module (i.e., the suction tube) was oriented directly toward the odor source (0°) compared to an orthogonal orientation (two-tailed t-test; p < 0.05). This directionality arises because the active sniffing module restricts air intake to the frontal sector, effectively creating a defined ’field of view’ that enables orientation based on signal amplitudes.

### 3.3 Odor Source Localization with Sniffbot: The Trident Algorithm

Next, we sought to test Sniffbot’s ability to localize an odor source in a windless (indoor) environment (**Fig 1C** and **Figure. 3A&B**). Each localization experiment started with Sniffbot positioned at a designated starting point of a 3.8-m × 4.8-m rectangular arena (**Figure. 3A & B**). The robot’s task was to use an odor-localization algorithm to autonomously reach a target square at the far end of the arena (40 cm × 40 cm, approximately twice the length of Sniffbot), where an odorant source was positioned (**Movie 2**). Success was defined as an arrival of the robot into the square within the predefined number of steps allowed per trial (e.g., 50, see **Materials and Methods**). During each experiment, Sniffbot stayed within the walls of the arena by utilizing its ultrasonic-based obstacle avoidance capacity.

Localization is a two-part problem including first odor detection and then movement towards the odor source. We tested three navigation algorithms [39]. The first two are established algorithms for odorant localization: E. coli (random walk) (**Figure. 3C**) and Spiral (**Figure. 3D**). The third, Trident, was developed as part of this study, based on the classical Hex-path paradigm (see **Supplementary Information**). These algorithms were modified to operate on a binary input—odor detected / no odor detected—where a single detected peak is sufficient to indicate odor detection (see **Supplementary Information**). Detection was defined as a signal with an amplitude exceeding a predetermined threshold, set individually for each antenna based on its ambient air EAG baseline, to account for sensitivity differences among antennae. Detailed descriptions of the algorithms’ logic are provided in the SI data; we briefly summarize them in what follows.

***E. coli***: Upon detecting an odor, the robot moves forward m units; otherwise, it turns randomly (between 0-360°) and moves forward n units (n<m, **Figure. 3C, Movie 3**).

***Spiral:*** Upon detecting an odor, the robot moves forward; otherwise, it follows a spiral movement pattern until it detects an odor (90° turns for up to 6 steps, **Figure. 3D, Movie 4**).

***Trident:*** The robot first samples 60° to one side: if it detects an odor, it turns and moves in that direction; if not, it samples 60° to the opposite side (120° apart) and again moves upon odor detection. If both sides are negative, it proceeds forward (**Figure. 3E, Movie 2**).

Representative trajectories from both successful and failed experiments using each of the different algorithms are presented in **Figure. 3F–K** (successes: **Figure. 3F, G, & H**; failures: **Figure. 3I, J, & K** for failures; see **Supp. Figure. 4** for trajectory analysis).

We compared each of the three search algorithms to two different random-navigation control algorithms. The first was a “no events” control [39,40] in which Sniffbot followed the movement pattern dictated by the focal search algorithm for no odor detected (repeated ‘0’ inputs, e.g., **Supp. Figure. 5C**). Since this condition is completely devoid of odor stimuli—and thus the robot’s movements are not comparable to a trajectory that includes the movement pattern that follows odor detection—we also employed a “shuffled events” control. In this control, we randomly redistributed the positive and negative odor detection events observed during odor-guided trials (i.e., the ‘0’ and the ‘1’ inputs), thereby preserving the overall movement pattern, yet with no relevance or direct influence of the odor source location (e.g. **Supp. Figure. 5D**, see **Supplementary Information**).

**Figure 5:**
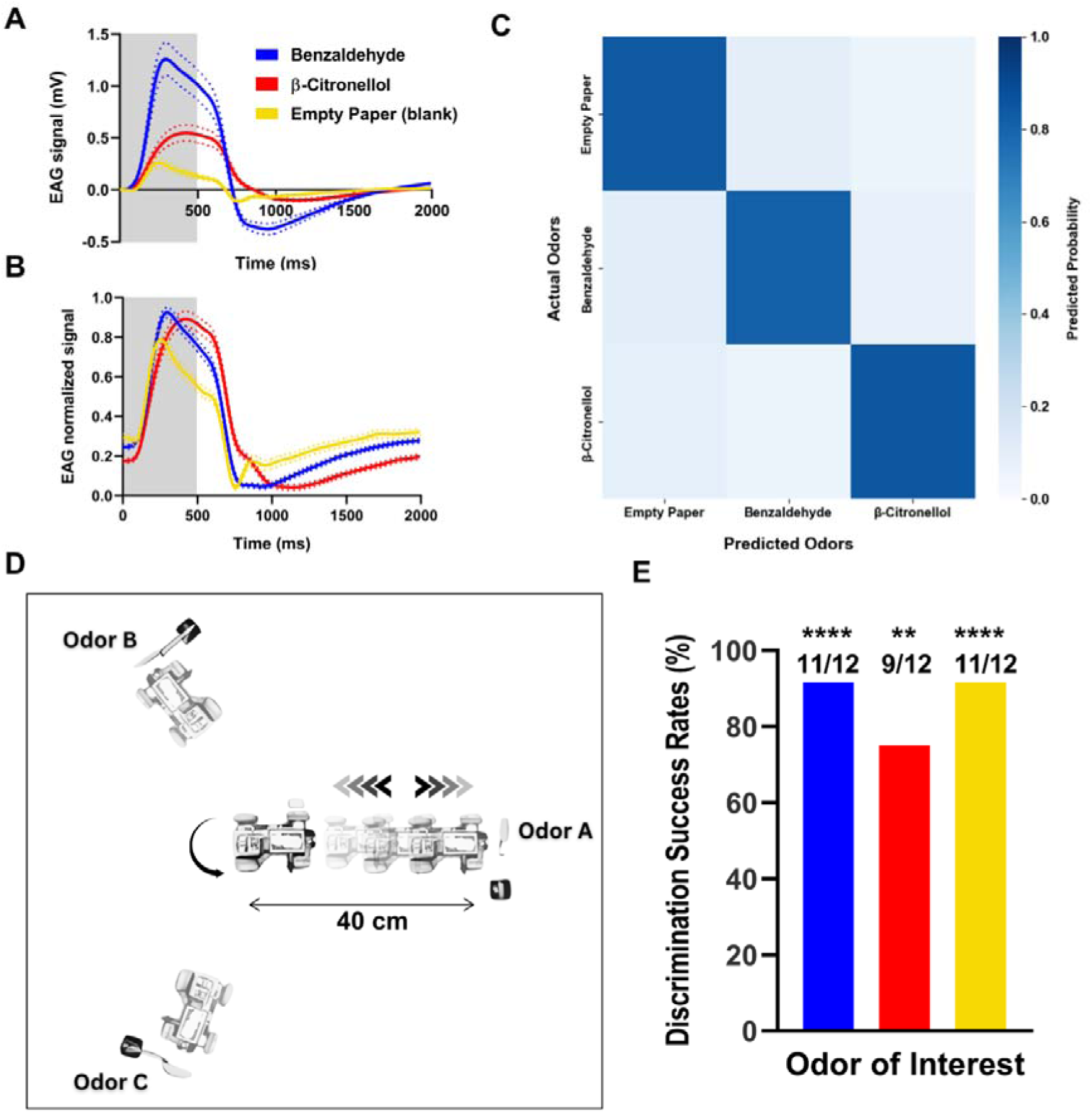
Sniffbot’s Odor Discrimination Capabilities. (A&B) Averaged and averaged-normalized EAG signals, respectively, of the training data set. The stimulus is represented by the gray square. N = 36 antennae, 318 repetitions. Dashed lines indicate the standard error. **(C)** Confusion matrix for the discrimination training dataset (normalized odorant responses as shown in panel (B)) using the logistic regression classifier, with an average accuracy of 83.33% over 318 samples. Two-tailed binomial test: p < 0.0001. **(D)** Schematic of the experimental procedure. **(E)** Discrimination success rates. N = 12 antennae, for each odor of interest. Two-tailed binomial test: ****, p < 0.0001; **, p < 0.01.

As shown in **Figure. 3L-N**, with a 50-step limit, all three odor-guided searching algorithms significantly outperformed each of the random navigation controls in terms of success rate (i.e., whether or not they reached the target square). When comparing the three navigation algorithms, the Trident’s success rate was highest (>90%), followed by the Spiral algorithm (slightly over 50%), while the E. coli algorithm was the least successful (just over 30%) (**Figure. 3O**); the Trident significantly outperformed the other two (two-tailed binomial test; p < 0.01). For the E. coli algorithm, success rates were initially low (∼35%). To test its robustness, we doubled its step limit to 100, which increased the success rate to ∼70%, while still outperforming the random navigation strategies (**Supp. Figure. 5A**, p < 0.0001). Conversely, we reduced the step limit for the best-performing Trident algorithm to 25 steps. Even under this constraint, Trident continued to outperform the random strategies (**Supp. Figure. 5B**).

Among the successful trials in which Sniffbot found the target, the Spiral and Trident algorithms did not significantly differ in either the number of steps required to reach the goal or the duration of the successful trials (step durations differed among algorithms, making the two comparisons non-equivalent. **Fig 3P & Q)**. The E. coli algorithm was significantly less efficient, requiring more steps and more time to reach the target compared with either of the other two algorithms (**Fig 3P & Q,** two-tailed t-test between Spiral and E.coli: p < 0.05. Welch’s test between E. coli and Trident: p < 0.0005; see **Supplementary Information**).

### 3.4 Simulating the Odor Localization Algorithms

We subsequently used simulations to gain a more in-depth understanding of the performance of the various search algorithms. To this end, we first constructed a map of odor dispersal in our windless arena, by combining the experimental data from 63 trials and calculating the probability of Sniffbot to encounter the odor in each area (**Figure. 4A&B,** see **Supplementary Information**). Next, we simulated each of the search algorithms in the odor map we had generated (n=1000 simulations per algorithm, see **Materials and Methods**). These simulations confirmed that the Trident algorithm performs best in terms of localization success, while the E. coli algorithm performs the worst (**Figure. 4C**, two-tailed binomial test; p < 0.0001 for all comparisons).

Overall, the success rates in the simulations were significantly lower than those observed in the field experiments: 16.8% vs. 30% for the E. coli algorithm, 31% vs. 53% for Spiral, and 52.7% vs. 93% for Trident. The efficiency of simulations (i.e., steps to success) was only slightly lower than our experimental data: the Spiral algorithm was the least efficient in the simulation, while the Trident algorithm reached the goal in the fewest steps (**Figure. 4D**, two-tailed t-test; p < 0.01 between E. coli and Spiral and p < 0.0001 between Trident and the other two). Examples of successful trajectories for the simulated Sniffbot are shown in **Figure. 4E, 4G, and 4I** as well as in **Movies 5-7**, and examples of failed trajectories are presented in **Figure. 4F, 4H, and 4J** as well as in **Movies 8-10,** for the E. coli, Spiral, and Trident algorithms, respectively.

### 3.5 Utilizing Sniffbot for Odor Discrimination

After validating Sniffbot’s ability to autonomously identify and localize an odor source in an indoor environment, we tested its ability to discriminate between different odorants in the same environment (**Fig 1D**). To do so, we used a machine learning-based model trained on (more than 300) EAG signals of two odors (Benzaldehyde, β-citronellol) and a blank control (**Fig 5A** and see **Supplementary Information**). The signals were normalized by maximum amplitude, to eliminate classifier bias caused by amplitude differences between odorants and by varying antenna sensitivities.

A logistic regression (LR) classifier performed better than three other models tested (see **Supplementary Information**) in discriminating between the odors, achieving a balanced accuracy of 83.3%—significantly exceeding chance (33.3%, estimated by training a classifier on shuffled data, *p* < 0.0001, **Figure. 5B & C, Supp. Figure. 6D-G**).

Next, we integrated the machine learning model into Sniffbot and tested its performance under real-world conditions. In each trial, three samples (the two odorants and a blank control) were randomly placed in three fixed positions in the room. Sniffbot was programmed to approach each position and determine which of the three was the predetermined target odor of interest (OOI), using the pretrained LR model (**Figure. 5D, Supp. Figure. 6I, Movie. 11** and see **Materials and Methods**). As the data in **Figure. 5E** show, Sniffbot was able to correctly identify two of the odorants (Benzaldehyde and blank paper) in over 90% of the trials and the third (β-citronellol) in 75% of the trials, all at rates significantly higher than random chance (33.33%, two-tailed binomial test; p < 0.0001 and p < 0.01, respectively). Although the success rate for β-citronellol was the lowest, its probability scores (which are an output of the LR classifier) were the highest, indicating that when Sniffbot did identify the β-citronellol odor, it did so with high certainty **(Supp. Figure. 6H**, two-tailed t-test; p < 0.05).

## 4. Discussion

In this work, we presented a biohybrid sniffing and sensing robot that can autonomously detect, discriminate, and localize different odorants in real time in a windless environment. Sniffbot integrates three main modules: (i) a sensing module comprising an insect olfactory sensing organ and a miniaturized electrophysiology system; (ii) a sniffing module, which actively samples the air in the environment; and (iii) a decision-making module, which analyzes the sensory input in real time to either navigates toward the odor source or identify specific odors.

We used a locust antenna as our olfactory sensor. Locusts are easy to breed and handle. They offer a well-studied and well-described olfactory system that responds to a large variety of chemical odorants [52,54–58], including explosives [13], and even those specific to certain types of cancer cells [15]. In a recent study by Shvil et al. [9], we introduced and extensively investigated the biohybrid approach, focusing on the sensory performance of the locust antenna, such as specificity and sensitivity. That work demonstrated that electroantennogram (EAG) signals recorded from the locust antenna can be successfully used to identify a large number of odorants presented individually, across different concentrations, as well as in mixtures. Here, we show that the locust antenna exhibits fast temporal dynamics, combining rapid response times (<1 s peak width) with recovery from habituation within tens of seconds (∼30 s). This high-temporal resolution contrasts with the slow recovery typical of conventional chemical sensors which often require minutes between measurements [59,60]. As a result, these temporal properties support Sniffbot’s theoretical high measurement throughput (>2,000 measurements per day) compared with analytical techniques such as GC–MS, which typically require 20-100 minutes per sample [61,62]. These properties allow rapid and continuous odor localization without the need for prolonged sensor resetting periods. The sensing module we developed for Sniffbot platform is a miniaturized, mobile version of the system used in Shvil et al. [9], with several necessary (though challenging) adaptations—including adjustments to the peak detection and analysis methodologies, improvement of the SNR, and introduction of a sniffing module. Notably, this measurement approach is substantially more straightforward and consistent than other methods of recording biological responses to odorants (e.g., intracellular recordings from olfactory brain centers [58]).

Our sensor’s success in discerning several odorants, using a machine-learning model, suggests that Sniffbot can be trained to detect any odorant that the locust’s antenna can perceive [9]. This capability is a substantial advantage over other reported robotic odorant detection systems. While pioneering bio-hybrid works have successfully utilized detached moth antennae [21,63,64] or Xenopus oocytes expressing insect receptors [65], these systems typically rely on specific ligand-receptor interactions (e.g., ethanol using electronical sensors, or female moth pheromone in bio-robots using moth antennae as sensors [18,52,63,66]). In contrast, our approach allows for the detection and classification of diverse odorants beyond a single target molecule. In this work, we used three odorants confirmed in our previous study to be detectable by the locust’s antenna. Specifically, we chose lemon oil for localization studies as it evokes a consistent response and, being naturally sourced, is safe for indoor use.

A key component of our Sniffbot is its sniffing module, which creates a timed air flow around the antenna to actively sample the surrounding environment. Active sniffing in the animal kingdom refers to the behavior in which mammals deliberately inhale air to enhance sensory perception and gather olfactory information from their environment [67–69]. Our custom-built sniffing apparatus, inspired by this behavior, enabled us to increase Sniffbot’s sensitivity (by increasing the concentration of odorant molecules in the vicinity of the sensor) and to reduce habituation by creating a transient (vs. constant or lingering) stimulus. The use of a timed airflow, followed by a designated recovery period, also enabled Sniffbot to sense multiple odorants consecutively without the residual effect of a previous odor. However, perhaps the most important feature is its ability to sample the air surrounding the robot without the need for external wind, therefore enabling localization in an indoor windless environment.

Indeed, localizing an odor source in an indoor windless environment is still considered a great challenge [40]. Unlike light or sound, whose dispersion can be readily measured [70], odor dispersal is extremely complex due to its reliance on multiple dynamic variables, making it very difficult to predict [37]. Moreover, recent research indicates that odors in natural environments, even when carried down-wind, do not disperse uniformly; instead, they form “odor patches" and local vortices [31,71,72]. To date, experiments involving odor source localization using bio-hybrid robots have been performed exclusively in settings with a well-defined directional wind source. In these setups, the odor source is strategically placed up-wind from the robot, ensuring that the wind directs the volatile compounds toward the sensor. Consequently, the robot simply needs to position itself against the wind and advance up-wind when it detects the odor. These experiments have given rise to the effective, bio-inspired "cast and surge" strategy, where the robot uses a zigzag pattern perpendicular to the wind (cast), and whenever it senses an odor, it moves directly upwind toward the source (surge) [31]. However, if the odor is not aligned with the wind or if there is no wind at all, these strategies fail. More complex strategies such as infotaxis or reinforcement learning, while powerful, also typically assume continuous concentration measurements or wind information. Odor source localization in indoor, windless environments is relatively underexplored, and in the limited literature available, researchers generally rely on one of three main algorithms: the E. coli and Spiral, both considered in this paper, and the Hex-Path algorithm, which served as the conceptual foundation for the novel Trident algorithm [34].

Although not the primary goal of this study, Trident—a novel navigation strategy developed in the course of our experiments—significantly outperformed both the *E. coli* and Spiral algorithms in terms of success rate, and also outperformed *E. coli* in the number of steps and time required to localize a target odorant source. We suggest that Trident’s success can be partially attributed to its movement pattern in response to a negative-input. The benefit of this movement pattern, which entails crossing the arena in straight lines is evidenced by the high success rates in the “no-event control” scenario. This motion pattern enables Sniffbot to efficiently sample its surroundings without methodically covering every point. In contrast, a point-by-point sampling approach such as that used by E. coli often leads to unnecessary, time-consuming sampling events that delay source localization. Although the movement pattern of Trident is more efficient, the two sides sampling protocol in each point leads to absolute localization times of 250–1700 s. This reflects the inherently challenging nature of plume-less environments where odor disperses via random pockets rather than informative gradients, necessitating extensive area coverage. A further optimization of this algorithm might be needed to shorten the localization times.

We note that the algorithms presented in this paper do not rely on memory of previous inputs when making decisions. Taking previous detection trends into account, for example, assigning greater weight to positive detections that follow other positives—could potentially enhance performance in future implementations.

To generate a more detailed comparison of the algorithms, we used the data from our experiments to create an odor concentration map and subsequently simulated the different algorithms’ performance in this environment. The simulations produced trends similar to those observed in our experiments regarding the algorithms’ relative success probability. However, the simulated success rates were approximately 1.8 times lower than those observed in the real-world experiments. One possible explanation for this discrepancy is the inaccurate estimation of the robot’s likelihood of encountering an odor at each point in space. In our simulation, the robot received a binary input (either positive or negative for odor detection) based solely on the probability measured in each grid cell—derived from the bio-hybrid robot’s odor measurements—with the assumption that each measurement was independent of the others. In reality, however, the presence of an odor at a given location is influenced by its detection at a neighboring point, as odor patches might extend across multiple grid cells. This interdependence means that the assumption of independent probabilities in the simulation could lead to inaccuracies, ultimately resulting in lower success rates than those observed in our field experiments.

One salient limitation of Sniffbot is the antenna’s limited lifespan. Though an insect antenna has recently been reported to remain vital for up to two weeks [73] when dissected off the animal, the locust antenna viable functionality in our system was limited to about 11 hours [9]. During this time, the EAG signal intensity declined exponentially, reaching 50% within the first hour of detachment. This issue could be mitigated by pre-calculating decay parameters and adjusting the odor detection function accordingly (e.g., by using a changing threshold). In this work, our experiments were conducted immediately after dissection, and we ensured that they did not exceed 45 minutes, making such adjustments unnecessary. Furthermore, because the sensor detects odors in a binary way, the signal’s amplitude decline is less important.

Our findings point to several promising avenues for future development. First, given our previous work showing that locust antennae can identify a large repertoire of very different odorants [9], an important next step is to assess the scope of Sniffbot’s identification and discrimination capabilities. Second, we tested our sensory system in windless indoor environments—a challenging and scientifically important context [40]. Future work could deploy Sniffbot in other environments—whether manmade or natural—where air currents carry odorants over greater distances; Such air currents could provide additional directional cues, potentially enhancing the sensor’s odor-detection capabilities and enabling the implementation of other anemotaxis-based algorithms. In addition, we implemented Sniffbot with ultrasonic sensors, which enabled the robotic vehicle to avoid collisions with the arena’s walls. The system could be further augmented with other sensors—it can currently also accommodate infrared sensors and a camera—for enhanced sensory input and potential improvement of navigation. Last, applications integrating multiple Sniffbots, either working as a robotic team (with dedicated roles) or taking advantage of swarm wisdom [47], could push the boundaries of the technology much further.

## Acknowledgments

General

We would like to thank Eliran Farhi, Peleg Levin, Itamar Mishani, and Alon Mizrahi for laying the foundations for the recording code and machine learning models. Additionally, we thank the staff of the Zoology Department for allowing us to place our arena in the seminar room for the navigation experiments. Special thanks to Roi Peled for providing the carbon air filter for the second version of the sniffer, to Elad Ozeri for modeling Sniffbot in SolidWorks and to Alexey Chizhik for the art work.

## ***4.1*** Author contributions

Conceptualization, Data Curation, Formal Analysis, Investigation, Methodology, Project Administration, Software, Visualization, Writing – Original Draft, Writing – Review & Editing: NS Software: NG Methodology: AN Software: OY Conceptualization, Funding Acquisition, Project Administration, Supervision, Visualization, Writing – Review & Editing: YY Conceptualization, Funding Acquisition, Project Administration, Supervision, Visualization, Writing – Review & Editing: BMM Conceptualization, Funding Acquisition, Project Administration, Supervision, Visualization, Writing – Review & Editing: AA

## 4.2 Competing interests

The authors declare that they do not have any conflict of interest.

## 5. Data and materials availability

All the data is available upon request.

Received: ((will be filled in by the editorial staff)) Revised: ((will be filled in by the editorial staff)) Published online: ((will be filled in by the editorial staff))

## Funding

Israeli Ministry of Science, Technology and Space MAFAT.

## Supporting Information

***Supplementary materials and methods*** Average Peaks’ Amplitude and Normalization Development of the Trident Navigation Algorithm Algorithms’ Logic – Pseudo Code

Navigation Experiments: Comparing Control Conditions and Accessing Algorithms Efficiency Odor Dispersal Mapping

Improved version of the sniffer

Machine Learning Model Choice and Training

## Supplementary figures

Figure. S1: Sniffbot’s components.

Figure. S2: Sniffing module, and distance and orientation discrimination. Figure. S3: Peak detection procedure.

Figure. S4: Robot trajectory analysis and plotting.

Figure. S5: Sniffbot’s odor localization experiments and controls. Figure. S6: Sniffbot’s odor discrimination experiment.

## Supplementary Movies

Movie. S1: Sniffbot preparation and experimental procedure.

Movie. S2: Sniffbot’s navigation experiment using the Trident algorithm, fast-forwarded ×10. Movie. S3: Example trajectory using the E.Coli algorithm: blue points indicate odor detection; red points indicate no detection (left panel); corresponding EAG signal at each point shown on the right panel. Peak detection parameters are displayed on the graph.

Movie S4: Same as Movie 3, using the Spiral algorithm.

Movie S5: Simulated Sniffbot’s navigation in a virtual odor dispersal map using the E.Coli algorithm (successful trial).

Movie S6: Same as Movie 5, using the Spiral algorithm. Movie S7: Same as Movie 5, using the Trident algorithm.

Movie. S8: Simulated Sniffbot’s navigation in a virtual odor dispersal map using the E.Coli algorithm (failed trial).

Movie. S9: Same as Movie 8 using the Spiral algorithm. Movie. S10: Same as Movie 8 using the Trident algorithm.

Movie. S11: Sniffbot’s discrimination experiment with corresponding recorded EAG signals.

*References cited only in the supplementary materials section: [74–81]

## Notes

### Competing Interest Statement

The authors have declared no competing interest.

## References

[1] T. Wasilewski, J. Gębicki, W. Kamysz, Bio-inspired approaches for explosives detection, TrAC - Trends Anal. Chem. 142 (2021) 116330. 10.1016/j.trac.2021.116330.

[2] J. Schuberth, Volatile organic compounds determined in pharmaceutical products by full evaporation technique and capillary gas chromatography/ ion-trap detection, Anal. Chem. 68 (1996) 1317–1320. 10.1021/ac951084a.

[3] Y. Xu, W. Cheung, C.L. Winder, R. Goodacre, VOC-based metabolic profiling for food spoilage detection with the application to detecting Salmonella typhimurium-contaminated pork, Anal. Bioanal. Chem. 397 (2010) 2439–2449. 10.1007/S00216-010-3771-Z.

[4] P.S. Murvay, I. Silea, A survey on gas leak detection and localization techniques, J. Loss Prev. Process Ind. 25 (2012) 966–973. 10.1016/J.JLP.2012.05.010.

[5] S. Rahman, A.S. Alwadie, M. Irfan, R. Nawaz, M. Raza, E. Javed, M. Awais, Wireless e-nose sensors to detect volatile organic gases through multivariate analysis, Micromachines 11 (2020) 597. 10.3390/MI11060597.

[6] H. Michelot, S. Chadwick, M. Morelato, M. Tahtouh, C. Roux, The screening of identity documents at borders for forensic drug intelligence purpose, Forensic Chem. 18 (2020) 100228. 10.1016/j.forc.2020.100228.

[7] D. Duff, C. Lennard, Y. Li, C. Doyle, K.J. Edge, I. Holland, K. Lothridge, P. Johnstone, P. Beylerian, V. Spikmans, Portable gas chromatography–mass spectrometry method for the in-field screening of organic pollutants in soil and water at pollution incidents, Environ. Sci. Pollut. Res. Int. 30 (2023) 93088. 10.1007/S11356-023-28648-W.

[8] A. Marcillo, J.C. Baca Cabrera, A. Widdig, C. Birkemeyer, A comparison between mobile and stationary gas chromatography–mass spectrometry devices for analysis of complex volatile profiles, Anal. Bioanal. Chem. 415 (2023) 137–155. 10.1007/S00216-022-04391-Y.

[9] N. Shvil, A. Golan, Y. Yovel, A. Ayali, B.M. Maoz, The locust antenna as an odor discriminator, Biosens. Bioelectron. 221 (2023) 114919. 10.1016/j.bios.2022.114919.

[10] Y. Li, X. Wei, Y. Zhou, J. Wang, R. You, Research progress of electronic nose technology in exhaled breath disease analysis, Microsystems Nanoeng. 9 (2023) 1–22. 10.1038/S41378-023-00594-0.

[11] L.R. Burnett, N.R. Hebdon, P.A. Stevens, M.D. Moljo, L.D. Waldrop, L.E. DeGreeff, Dog sniffing biomechanic responses in an odor detection test of odorants with differing physical properties, J. Anim. Sci. 102 (2024) skae353. 10.1093/JAS/SKAE353.

[12] L.S. Fernandez, S.A. Kane, M.T. DeChant, P.A. Prada-Tiedemann, N.J. Hall, Environmental effects on explosive detection threshold of domestic dogs, PLoS One 19 (2024) e0306817. 10.1371/journal.pone.0306817.

[13] D. Saha, D. Mehta, E. Altan, R. Chandak, M. Traner, R. Lo, P. Gupta, S. Singamaneni, S. Chakrabartty, B. Raman, Explosive sensing with insect-based biorobots, Biosens. Bioelectron. X 6 (2020) 100050. 10.1016/j.biosx.2020.100050.

[14] D. Romano, E. Donati, G. Benelli, C. Stefanini, A review on animal–robot interaction: from bio-hybrid organisms to mixed societies, Biol. Cybern. 113 (2019) 201–225. 10.1007/s00422-018-0787-5.

[15] A. Farnum, M. Parnas, E. Hoque Apu, E. Cox, N. Lefevre, C.H. Contag, D. Saha, Harnessing insect olfactory neural circuits for detecting and discriminating human cancers, Biosens. Bioelectron. 219 (2023) 114814. 10.1016/j.bios.2022.114814.

[16] K.-E. Kaissling, Responses of Insect Olfactory Neurons to Single Pheromone Molecules, Olfactory Concepts Insect Control Alternative to Insectic. (2019) 1–27. 10.1007/978-3-030-05165-5_1.

[17] M. Schott, C. Wehrenfennig, T. Gasch, A. Vilcinskas, Insect antenna-based biosensors for in situ detection of volatiles, Adv. Biochem. Eng. Biotechnol. 136 (2013) 101–122. 10.1007/10_2013_210.

[18] F. Van Breugel, J. Riffell, A. Fairhall, M.H. Dickinson, Mosquitoes use vision to associate odor plumes with thermal targets, Curr. Biol. 25 (2015) 2123–2129. 10.1016/j.cub.2015.06.046.

[19] J.S. Elkinton, C. Schal, T. Ono, R.T. Cardé, Pheromone puff trajectory and upwind flight of male gypsy moths in a forest, Physiol. Entomol. 12 (1987) 399–406. 10.1111/j.1365-3032.1987.tb00766.x.

[20] M. Ai, S. Min, Y. Grosjean, C. Leblanc, R. Bell, R. Benton, G.S.B. Suh, Acid sensing by the Drosophila olfactory system, Nature 468 (2010) 691–695. 10.1038/nature09537.

[21] D. Terutsuki, T. Uchida, C. Fukui, Y. Sukekawa, Y. Okamoto, R. Kanzaki, Real-time odor concentration and direction recognition for efficient odor source localization using a small bio-hybrid drone, Sensors Actuators, B Chem. 339 (2021) 129770. 10.1016/j.snb.2021.129770.

[22] C. Jiang, H. Xu, L. Yang, J. Liu, Y. Li, K. Takei, W. Xu, Neuromorphic antennal sensory system, Nat. Commun. 15 (2024) 2239. 10.1038/S41467-024-46393-7.

[23] I. Fishel, Y. Amit, N. Shvil, A. Sheinin, A. Ayali, Y. Yovel, B.M. Maoz, Ear-Bot: locust ear-on-a-chip bio-hybrid platform, Sensors 21 (2021) 228. 10.3390/S21010228.

[24] H.D. Nguyen, V.T. Dung, H. Sato, T.T. Vo-Doan, Efficient autonomous navigation for terrestrial insect-machine hybrid systems, Sensors Actuators B Chem. 376 (2023) 132988. 10.1016/J.SNB.2022.132988.

[25] K.C. Park, Odor discrimination using insect electroantennogram responses from an insect antennal array, Chem. Senses 27 (2002) 343–352. 10.1093/chemse/27.4.343.

[26] D.H. Slone, B.T. Sullivan, An automated approach to detecting signals in electroantennogram data, J. Chem. Ecol. 33 (2007) 1748–1762. 10.1007/S10886-007-9338-6.

[27] A.J. Myrick, K.C. Park, J.R. Hetling, T.C. Baker, Detection and discrimination of mixed odor strands in overlapping plumes using an insect-antenna-based chemosensor system, J. Chem. Ecol. 35 (2009) 118–130. 10.1007/s10886-008-9582-4.

[28] R.T. Cardé, M.A. Willis, Navigational strategies used by insects to find distant, wind-borne sources of odor, J. Chem. Ecol. 34 (2008) 854–866. 10.1007/S10886-008-9484-5.

[29] M.J. Anderson, J.G. Sullivan, T.K. Horiuchi, S.B. Fuller, T.L. Daniel, A bio-hybrid odor-guided autonomous palm-sized air vehicle, Bioinspir. Biomim. 16 (2021) 026002. 10.1088/1748-3190/ABBD81.

[30] T. Lazebnik, Y. Golov, R. Gurka, A. Harari, A. Liberzon, Exploration-exploitation model of moth-inspired olfactory navigation, J. R. Soc. Interface 21 (2024) 20230746. 10.1098/RSIF.2023.0746.

[31] S.P. Wechsler, V. Bhandawat, Behavioral algorithms and neural mechanisms underlying odor-modulated locomotion in insects, J. Exp. Biol. 226 (2023) jeb200261. 10.1242/JEB.200261.

[32] N.J. Vickers, Mechanisms of animal navigation in odor plumes, Biol. Bull. 198 (2000) 203–212. 10.2307/1542524.

[33] T. Emonet, M. Vergassola, Olfactory cues and memories in animal navigation, Nat. Rev. Phys. 6 (2024) 215–216. 10.1038/s42254-024-00710-7.

[34] X. Chen, J. Huang, Odor source localization algorithms on mobile robots: A review and future outlook, Rob. Auton. Syst. 112 (2019) 123–136. 10.1016/j.robot.2018.11.014.

[35] R. Gasser, Measurement of small air velocities in air-conditioned rooms, Air Infiltration and Ventilation Centre, 1986. https://www.aivc.org/resource/measurement-small-air-velocities-air-conditioned-rooms-location-europe (accessed June 12, 2025).

[36] Κ. Niachou, M. Santamouris, C. Georgakis, Natural and Hybrid Ventilation in the Urban Environment - Technical Note 61, 2007. https://www.aivc.org/resource/tn-61-natural-and-hybrid-ventilation-urban-environment.

[37] G. Reddy, V.N. Murthy, M. Vergassola, Olfactory sensing and navigation in turbulent environments, Annu. Rev. Condens. Matter Phys. 13 (2022) 191–213. 10.1146/annurev-conmatphys-031720-032754.

[38] G. Kowadlo, R.A. Russell, Robot odor localization: A taxonomy and survey, Int. J. Rob. Res. 27 (2008) 869–894. 10.1177/0278364908095118.

[39] Y. Yang, Q. Feng, H. Cai, J. Xu, F. Li, Z. Deng, C. Yan, X. Li, Experimental study on three single-robot active olfaction algorithms for locating contaminant sources in indoor environments with no strong airflow, Build. Environ. 155 (2019) 320–333. 10.1016/j.buildenv.2019.03.043.

[40] G. Ferri, E. Caselli, V. Mattoli, A. Mondini, B. Mazzolai, P. Dario, SPIRAL: A novel biologically-inspired algorithm for gas/odor source localization in an indoor environment with no strong airflow, Rob. Auton. Syst. 57 (2009) 393–402. 10.1016/j.robot.2008.07.004.

[41] R.A. Heinonen, L. Biferale, A. Celani, M. Vergassola, Optimal policies for Bayesian olfactory search in turbulent flows, Phys. Rev. E 107 (2023) 055105. 10.1103/PhysRevE.107.055105.

[42] M. Vergassola, E. Villermaux, B.I. Shraiman, ‘Infotaxis’ as a strategy for searching without gradients, Nature 445 (2007) 406–409. 10.1038/nature05464.

[43] R.A. Heinonen, L. Biferale, A. Celani, M. Vergassola, Exploring Bayesian olfactory search in realistic turbulent flows, Phys. Rev. Fluids 10 (2025) 64614. 10.1103/9q6q-nlxc.

[44] K.V.B. Verano, E. Panizon, A. Celani, Olfactory search with finite-state controllers, Proc. Natl. Acad. Sci. U. S. A. 120 (2023) e2304230120. 10.1073/PNAS.2304230120.

[45] N. Rigolli, G. Reddy, A. Seminara, M. Vergassola, Alternation emerges as a multi-modal strategy for turbulent odor navigation, Elife 11 (2022) e76989. 10.7554/eLife.76989.

[46] V. Jayaram, N. Kadakia, T. Emonet, Sensing complementary temporal features of odor signals enhances navigation of diverse turbulent plumes, Elife 11 (2022) e72415. 10.7554/eLife.72415.

[47] J. Wang, Y. Lin, R. Liu, J. Fu, Odor source localization of multi-robots with swarm intelligence algorithms: a review, Front. Neurorobot. 16 (2022) 949888. 10.3389/FNBOT.2022.949888.

[48] Y. Jia, S. Fan, W. Cui, C. Di, Y. Hao, A novel distributed hybrid cognitive strategy for odor source location in turbulent and sparse environment, Entropy 27 (2025) 826. 10.3390/E27080826.

[49] W. Takken, B.G.J. Knols, Olfaction in vector-host interactions, Wageningen Academic Publishers, 2010. 10.3920/978-90-8686-698-4.

[50] D. Wicher, Tuning insect odorant receptors, Front. Cell. Neurosci. 12 (2018) 94. 10.3389/FNCEL.2018.00094.

[51] S. Nizampatnam, D. Saha, R. Chandak, B. Raman, Dynamic contrast enhancement and flexible odor codes, Nat. Commun. 9 (2018) 3055. 10.1038/s41467-018-05533-6.

[52] B. Raman, J. Joseph, J. Tang, M. Stopfer, Temporally diverse firing patterns in olfactory receptor neurons underlie spatiotemporal neural codes for odors, J. Neurosci. 30 (2010) 1994–2006. 10.1523/jneurosci.5639-09.2010.

[53] S. Haney, D. Saha, B. Raman, M. Bazhenov, Differential effects of adaptation on odor discrimination, J. Neurophysiol. 120 (2018) 171–185. 10.1152/jn.00389.2017.

[54] X. Jiang, H. Breer, P. Pregitzer, Sensilla-specific expression of odorant receptors in the desert locust Schistocerca gregaria, Front. Physiol. 10 (2019) 1052. 10.3389/FPHYS.2019.01052.

[55] S.A. Ochieng, E. Hallberg, B.S. Hansson, Fine structure and distribution of antennal sensilla of the desert locust, Schistocerca gregaria (Orthoptera: Acrididae), Cell Tissue Res. 291 (1998) 525–536. 10.1007/S004410051022.

[56] B.S. Hansson, S. Anton, Function and morphology of the antennal lobe: New developments, Annu. Rev. Entomol. 45 (2000) 203–231. 10.1146/annurev.ento.45.1.203.

[57] M. Stopfer, V. Jayaraman, G. Laurent, Intensity versus identity coding in an olfactory system, Neuron 39 (2003) 991–1004. 10.1016/j.neuron.2003.08.011.

[58] D. Saha, K. Leong, N. Katta, B. Raman, Multi-unit recording methods to characterize neural activity in the locust (Schistocerca americana) olfactory circuits, J. Vis. Exp. (2013) e50139. 10.3791/50139.

[59] X. Tang, D. Lahem, J.-P. Raskin, M. Debliquy, A hybrid gas sensor based on compound of graphene with polypyrrole, in: XXXI International Conference on Surface Modification Technologies (SMT31), 2016.

[60] G.F. Fine, L.M. Cavanagh, A. Afonja, R. Binions, Metal oxide semi-conductor gas sensors in environmental monitoring, Sensors 10 (2010) 5469–5502. 10.3390/S100605469.

[61] A.B. Fialkov, S.J. Lehotay, A. Amirav, Less than one minute low-pressure gas chromatography - mass spectrometry, J. Chromatogr. A 1612 (2020) 460691. 10.1016/j.chroma.2019.460691.

[62] R.A. Hites, Gas Chromatography Mass Spectrometry, in: F. Settle (Ed.), Handb. Instrum. Tech. Anal. Chem., Prentice Hall, Upper Saddle River, NJ, 1997: pp. 609–626.

[63] Y. Kuwana, S. Nagasawa, I. Shimoyama, R. Kanzaki, Synthesis of the pheromone-oriented behaviour of silkworm moths by a mobile robot with moth antennae as pheromone sensors, Biosens. Bioelectron. 14 (1999) 195–202. 10.1016/S0956-5663(98)00106-7.

[64] Y. Kuwana, I. Shimoyama, H. Miura, Steering control of a mobile robot using insect antennae, IEEE Int. Conf. Intell. Robot. Syst. 2 (1995) 530–535. 10.1109/IROS.1995.526267.

[65] N. Misawa, H. Mitsuno, R. Kanzaki, S. Takeuchi, Highly sensitive and selective odorant sensor using living cells expressing insect olfactory receptors, Proc. Natl. Acad. Sci. U. S. A. 107 (2010) 15340–15344. 10.1073/PNAS.1004334107.

[66] J. Horibe, N. Ando, R. Kanzaki, Odor-searching robot with insect-behavior-based olfactory sensor, Sensors Mater. 33 (2021) 4185–4202. 10.18494/SAM.2021.3369.

[67] M. Wachowiak, Active Sensing in Olfaction, in: A. Menini (Ed.), Neurobiol. Olfaction, CRC Press/Taylor & Francis, 2010: pp. 305–328. 10.1201/9781420071993-c12.

[68] M. Wachowiak, All in a sniff: olfaction as a model for active sensing, Neuron 71 (2011) 962–973. 10.1016/J.NEURON.2011.08.030.

[69] J. Crimaldi, H. Lei, A. Schaefer, M. Schmuker, B.H. Smith, A.C. True, J. V. Verhagen, J.D. Victor, Active sensing in a dynamic olfactory world, J. Comput. Neurosci. 50 (2022) 1–6. 10.1007/S10827-021-00798-1.

[70] S.M. Ross, Introduction to Probability Models, 10th ed., Academic Press, 2010.

[71] D. Martinez, L. Arhidi, E. Demondion, J.B. Masson, P. Lucas, Using insect electroantennogram sensors on autonomous robots for olfactory searches, J. Vis. Exp. (2014) e51704. 10.3791/51704.

[72] M. Pannunzi, T. Nowotny, Odor stimuli: not just chemical identity, Front. Physiol. 10 (2019) 1428. 10.3389/fphys.2019.01428.

[73] E.M. Steel, Z.E. Brooks, M. Kornexl, N. Vijai, M.A. Hawkins, A. Dixon, M. Willis, S.S. Kim, Biohybrid volatile organic compound sensing system, Proc. IEEE Natl. Aerosp. Electron. Conf. NAECON (2024) 251–256. 10.1109/NAECON61878.2024.10670636.

[74] J. Adler, The sensing of chemicals by bacteria, Sci. Am. 234 (1976) 40–47. 10.1038/scientificamerican0476-40.

[75] R.A. Russell, A. Bab-Hadiashar, R.L. Shepherd, G.G. Wallace, A comparison of reactive robot chemotaxis algorithms, Rob. Auton. Syst. 45 (2003) 83–97. 10.1016/S0921-8890(03)00120-9.

[76] R.A. Russell, Chemical source location and the Robomole project, Proc. Australas. Conf. Robot. Autom. (2003) 1–6.

[77] Y. Liu, X. Zhao, J. Xu, S. Zhu, D. Su, Rapid location technology of odor sources by multi-UAV, J. F. Robot. 39 (2022) 600–616. 10.1002/rob.22066.

[78] J. Burgués, V. Hernández, A.J. Lilienthal, S. Marco, Smelling nano aerial vehicle for gas source localization and mapping, Sensors 19 (2019) 478. 10.3390/s19030478.

[79] S. Shigaki, N. Minakawa, M. Yamada, H. Ohashi, D. Kurabayashi, K. Hosoda, Animal-in-the-loop system with multimodal virtual reality to elicit natural olfactory localization behavior, Sensors Mater. 33 (2021) 4211–4228. 10.18494/SAM.2021.3609.

[80] P. Ojeda, J. Monroy, J. Gonzalez-Jimenez, Robotic gas source localization with probabilistic mapping and online dispersion simulation, IEEE Trans. Robot. 40 (2024) 3551–3564. 10.1109/TRO.2024.3426368.

[81] V.H. Bennetts, A.J. Lilienthal, P.P. Neumann, M. Trincavelli, Mobile robots for localizing gas emission sources on landfill sites: Is bio-inspiration the way to go?, Front. Neuroeng. 4 (2012) 20. 10.3389/FNENG.2011.00020.

