## Supporting information for "The Sniffbot: A biohybrid robot for active sensing-based odor localization and discrimination"

### 1. **Materials and Methods**

***1.1 Average Peaks’ Amplitude and Normalization***

For the distance and directionality experiments (**Figure. 2G & H**), we calculated the average amplitude of all detected peaks within each recording to represent signal intensity. In the distance experiment (**Figure. 2G**), all values were further normalized to 7.626—the highest mean of the average peak amplitudes across recordings (**Supp. Figure. 2B**), corresponding to the sniffed odor at 50 cm. For the dose-response experiments (**Figure. 2F**), which included only one peak per recording, we used the peak’s maximum amplitude and normalized each concentration to the highest amplitude recorded for that specific antenna.

***1.2 Development of the Trident Navigation Algorithm***

The *Trident* navigation pattern was not initially designed as a primary contribution and was developed based on the classical Hex-path paradigm. In the classical Hex-path algorithm, the agent moves on a grid of hexagons, comparing the amplitudes of the present and previous signals [*34,39*]. When we modified this algorithm to operate on a binary input, Sniffbot entered an infinite loop in response to a consecutive number of “no odorant” inputs. This necessitated the creation of a modified algorithm. After observing its promising behavior empirically, we formalized the pattern into the *Trident* algorithm for controlled comparison and analysis.

***1.3 Algorithms’ Logic – Pseudo Code***

*E. coli Algorithm*

The E. coli algorithm is based on the movement of *E. coli* bacteria as first described by Adler (1976)[*74*] and later implemented in robotics by Russell et al. (2003)[*75]*. The adapted algorithm for Sniffbot is as follows **(Figure. 3C)**:

repeat {

if
 odor peak detected by Sniffbot then move forward *m* units (~40cm for Sniffbot) ;

else
 rotate in a random direction (±random(360°)) and move forward *n* units (where n < m, n=~20cm for Sniffbot);

}

*Note:* Additional variability is introduced by randomly choosing the turn direction (clockwise or counterclockwise).

*Spiral Algorithm:*
The Spiral algorithm, adapted from Yang et al. (2019)[*39*], was modified for Sniffbot as follows (**Figure. 3D**):

repeat {

if odor peak detected or the number of spiral steps > 6 then

move forward *m* units;

reset spiral step count;

choose a new spiral direction (6 steps);

else

turn in a spiral pattern and move forward *m* units;

increment spiral step count by one;

increase the step length by n;

}

*Note:* Variability is added by randomly choosing the spiral direction (clockwise or counterclockwise).

*Trident Algorithm:*
The Trident algorithm is a modification of the Hex-path algorithm described by Russell (2003)[*78*], adapted for Sniffbot **(Figure. 3E)**:

repeat
{

turn 60° to one side (chosen randomly) and sample;

if odor peak detected then

move forward *m* units;

else {

turn 120° to the opposite direction and sample;

if odor peak detected then

move forward *m* units;

else {

turn 60° to the opposite direction (returning to the original “straight” heading);
 move forward *m* units;

}

}

}

All algorithms are terminated if Sniffbot reaches the predetermined step capacity (usually 50 steps).

***1.4 Navigation Experiments: Comparing Control Conditions and Accessing Algorithms Efficiency***

Considering the performance of the two control conditions, we observed that, for both the E. coli and Spiral algorithms, the “shuffled events” control achieved higher success rates compared with the “no event” control (**Figure. 3L & M and Supp. Figure. 5A**). However, the Trident algorithm displayed the opposite effect. This difference is attributed to the fact that straight-line movement, characteristic of the negative-input response in the Trident algorithm, allows for more efficient ground coverage, thus increasing the chance of accidentally encountering the odor source (**Figure. 3N and Supp. Figure. 5B**).

When assessing the algorithm’s efficiency, we evaluated two parameters: the number of steps to reach the goal and the duration of the trial—considering only successful trials. While these two measures may appear redundant, both were necessary because, in the Trident algorithm, a single step can take twice as long when Sniffbot does not detect odor on the initially sampled side, prompting an additional sample before making a decision (**Figure. 3P & Q**).

***1.5 Odor Dispersal Mapping***

In windless environments, odorants diffuse to form a detectable gradient only within ~40 cm of the source [*31,37,38*], while beyond that range dispersal becomes complex, unpredictable, and challenging to measure. Hence, it is often approximated using a simulated Gaussian model [*77*]. Most of the existing odor dispersal maps, both experimental and simulated, are thus limited to odor dispersal in the presence of airflow [*31,37 ,78-81*]. The odor dispersal map presented herein is based on data from the experimental measurements. We mapped the arena used in our navigation experiments by dividing it into 20 × 20 cm squares and calculated the probability of encountering the odor in each, finally, the heatmap was smoothed with Gaussian smoothing (sigma factor = 4) **(Figure. 4A & B)**.

***1.6 Improved version of the sniffer***

To perform odor discrimination using Sniffbot, we further improved the active sensing system. The solenoid activation leads to a sudden burst of airflow on the antenna, which triggers a response from a population of neurons on the antenna equipped with tactile receptors **(Supp. Figure. 6A)**. Although this response decays within approximately 1–2 seconds, it initially masks the olfactory receptor neurons’ (ORNs’) response to the odor stimulus during the time period crucial for discrimination (the first 0.5 seconds). To address this issue, we added a custom-made carbon air filter (**Supp. Figure. 6B**), allowing a constant flow of filtered air to pass over the antenna, habituating the tactile neurons response. Upon activation of the "sniff", incoming surrounding air merges with the constant filtered airflow. This resulted in an approximately 50% reduction in the wind burst effect in terms of both amplitude and duration **(Supp. Figure. 6C),** and in a more robust olfactory response from the antenna**.**

***1.7 Machine Learning Model Choice and Training***

The optimal hyperparameters for each model were determined using the scikit-learn library in Python 3 (Python Software Foundation, Scotts Valley, CA) via the GridSearchCV module, followed by KFold cross-validation (due to the limited size of our training dataset). The chosen parameters for each model were as follows: (1) *Logistic regression* {'C': 10, 'max_iter': 100, 'penalty': 'l1', 'solver': 'liblinear', 'tol': 1e-05}, (2) *Random Forest* {'criterion': 'entropy', 'max_depth': 7, 'max_features': 'auto', 'n_estimators': 500}, (3) *Support Vector* *Classifier* (SVC) {'C': 1.0, 'degree': 3, 'kernel': 'poly', 'probability': True, 'tol': 0.001}, and (4) *k-nearest neighbors* (KNN) {'n_neighbors': 10, 'weights': 'distance'}.

The classifier's input consisted of the EAG signal that had been interpolated, filtered, and normalized (500 milliseconds of exposure within a 2000-millisecond recording period, normalized to its maximum value, **Fig 5A & B**). Among the tested models, the logistic regression classifier achieved the highest performance, with an area under curve (AUC) score of 0.957 and a balanced accuracy of 83.3% on the test data. To prevent data bias, we ensured that repetitions from the same antenna were not used in both the training and testing sets.

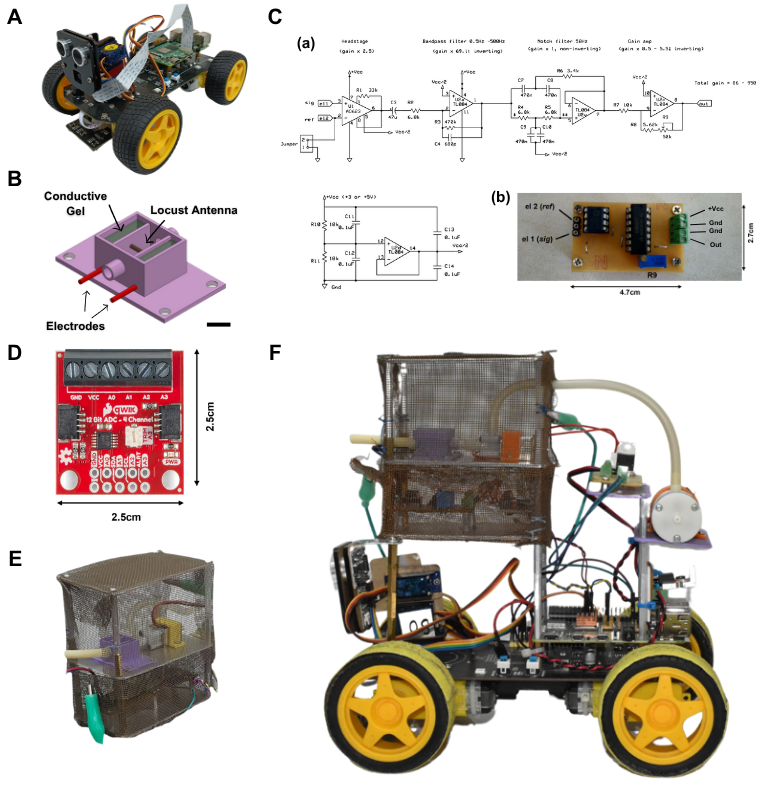
**Supplementary Figures:**

**Suppl. Figure. 1: Sniffbot’s Components.** (A) The robotic backbone, comprising the Freenove 4WD Smart Car Kit and Raspberry Pi® (B) A SolidWorks® sketch of the antenna holder with the antenna and electrodes (scale bar = 1 cm). (C) The custom-made EAG amplifier: (a) Circuit diagram and (b) Prototype. (D) The IDAC by Adafruit (ADS1015 device) (E) The custom-made Faraday cage. (C-E) constitute the miniaturized EAG recording system **(**F) A photograph of the assembled Sniffbot.

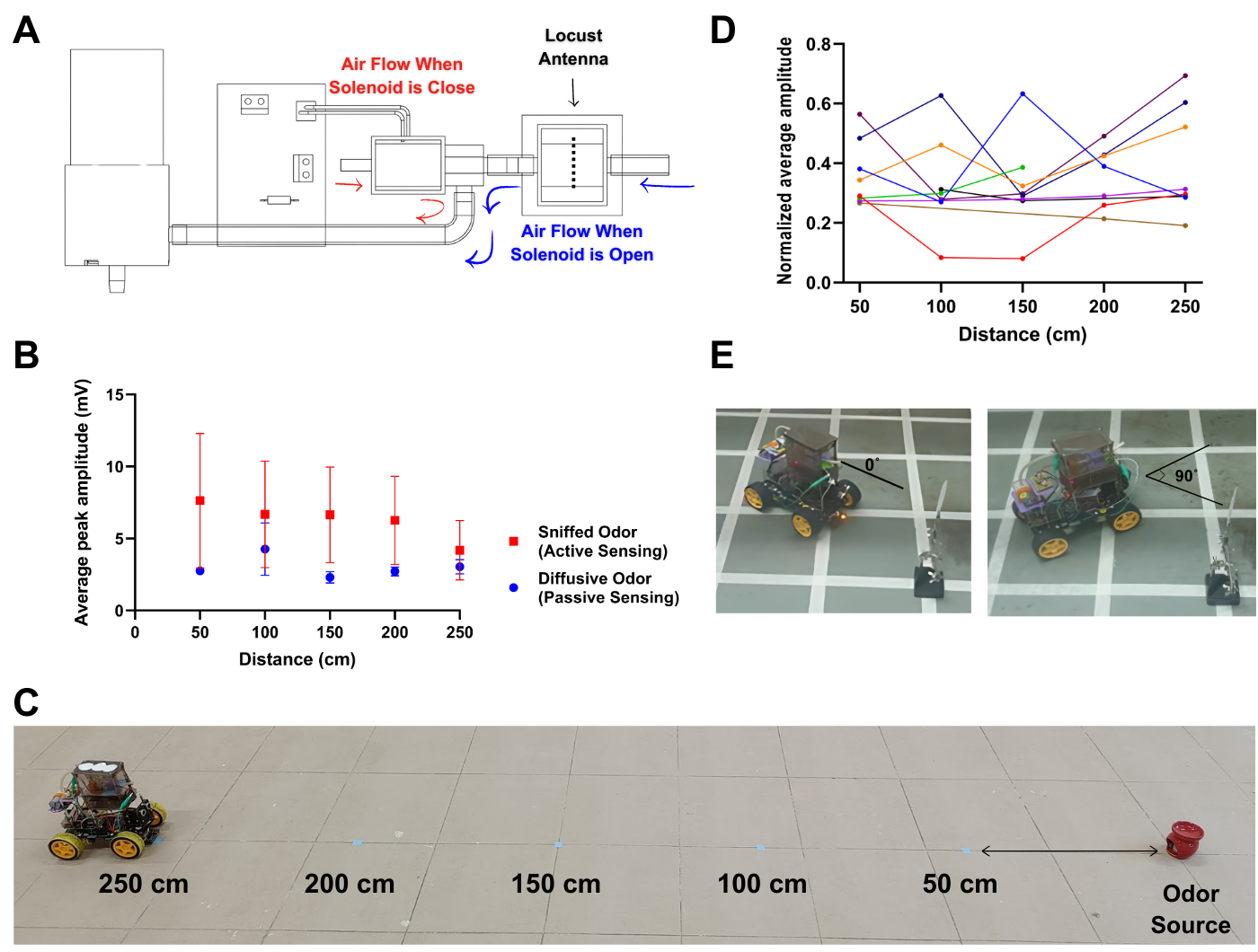

**Suppl. Figure. 2:** **Sniffing Module, and Distance and Orientation Discrimination.** (A) Sniffer air flow scheme: when the solenoid is closed, air circulates in a closed loop without passing through the antenna (red arrows). When the solenoid opens, the surrounding air is redirected to flow through the antenna (blue arrows). The locust antenna’s position is indicated by the black dashed line. (B) Average peak amplitude of the EAG signal as a function of Sniffbot’s distance from the odor source. Passive EAG shows no distance correlation (r=-0.2), confirming need for active sniffing in windless environments. N=10 antennae; 41 repetitions for passive experiments and N=9 antennae;25 repetitions for active experiments. Vertical lines represent standard error. Each recording lasts for 15 seconds (C) Layout of distance experiments (see *Figure 2H*). (D) Normalized mean EAG peak amplitude as a function of Sniffbot–source distance (Fig. 2G). Individual lines correspond to single antennae in the diffusive condition, demonstrating that the absence of correlation is not due to averaging. Only antennae with ≥3 measurements were included. (E). Sniffer directionality experiment layout. The robot was positioned 25 cm from the odor source.

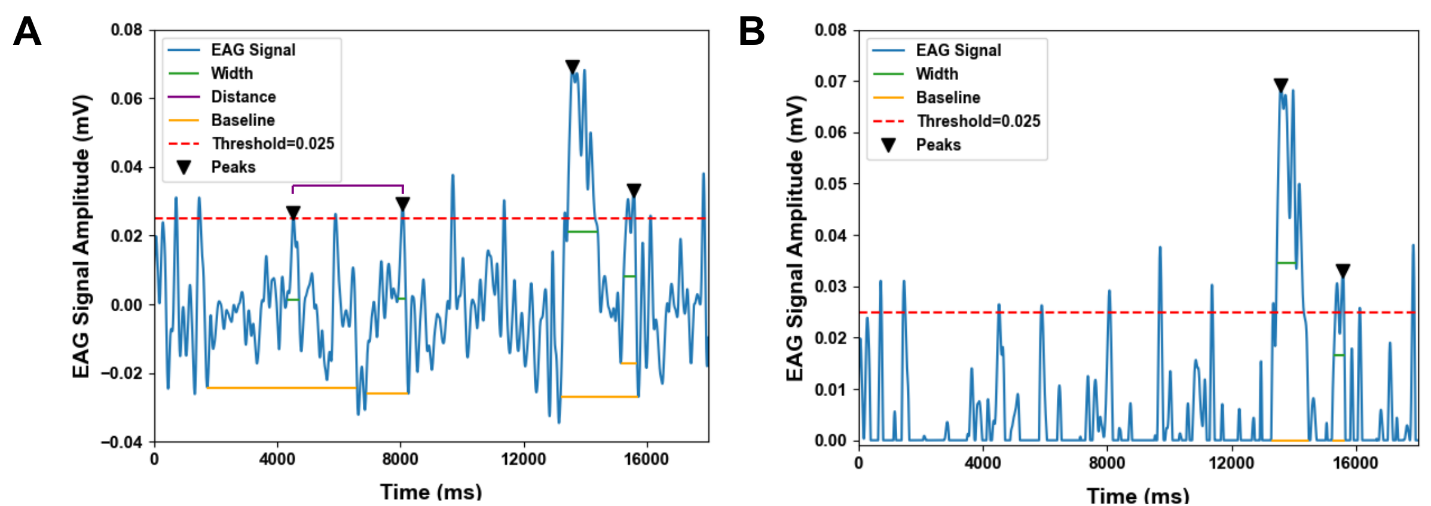

**Suppl. Figure. 3:** **Peak Detection Procedure.** (A) Peak detection paradigm for the entire signal and (B) for the positive fraction only. The automatically detected baseline is shown in orange, with the width at half-height indicated in green (relative height = 0.5). The minimum time interval between two peaks is indicated in purple (distance must be >1000ms), and detected peaks are marked with black triangles. The signal was recorded for 20 seconds.

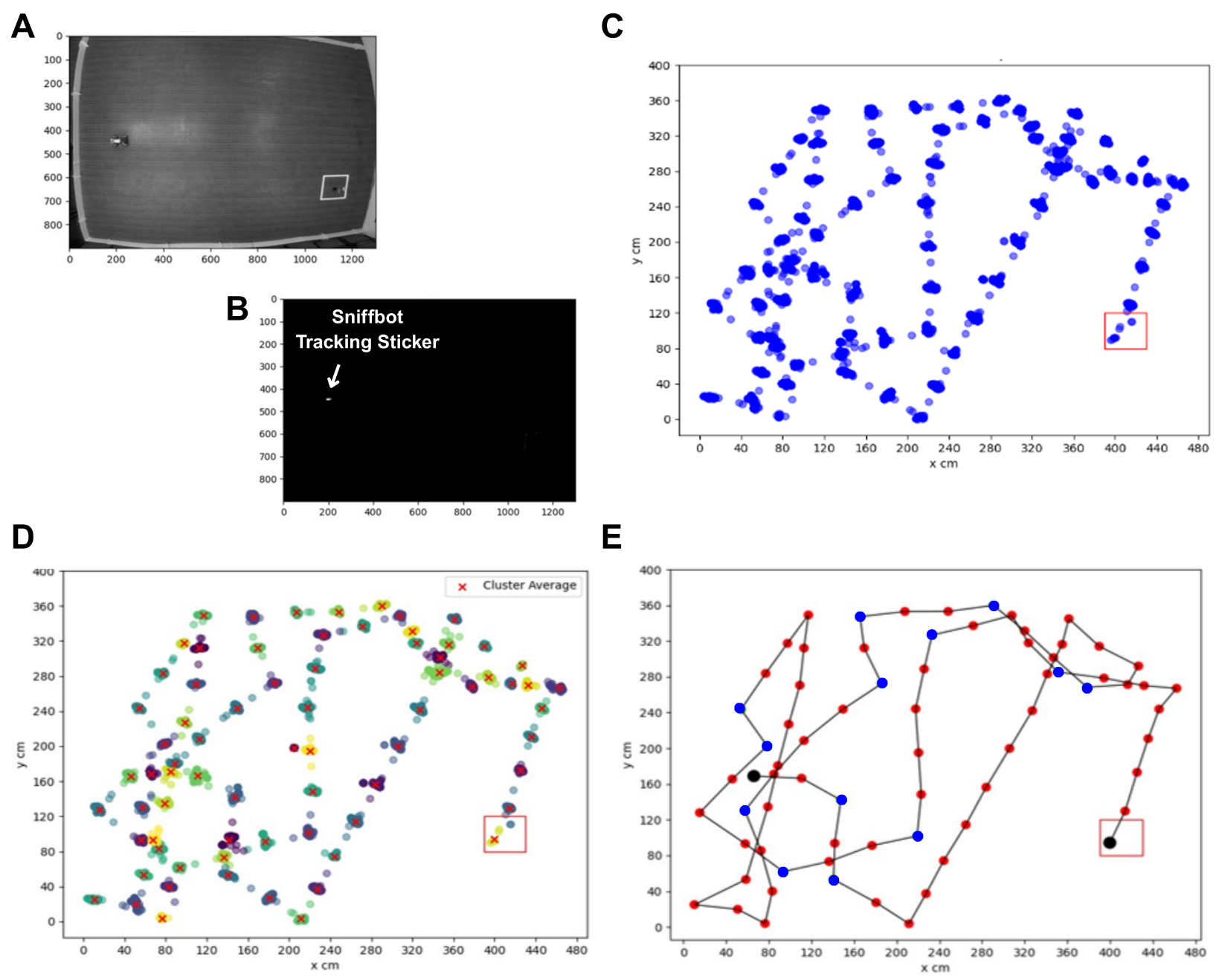

**Suppl. Figure. 4: Robot Trajectory Analysis and Plotting.** *Analysis Stage:* (A) Gray-shaded photograph of the experimental site. (B) High-contrast image highlighting only the white sticker on top of Sniffbot. *Plotting Stage*: (C) Scatter plot of all points in the room where Sniffbot passed, based on code detection of the white sticker. (D) Identification and plotting of the clusters’ average. (E) Color mapping of the clusters’ averages according to odor detection, with odor-detected areas shown in blue and areas with no odor detected shown in red.

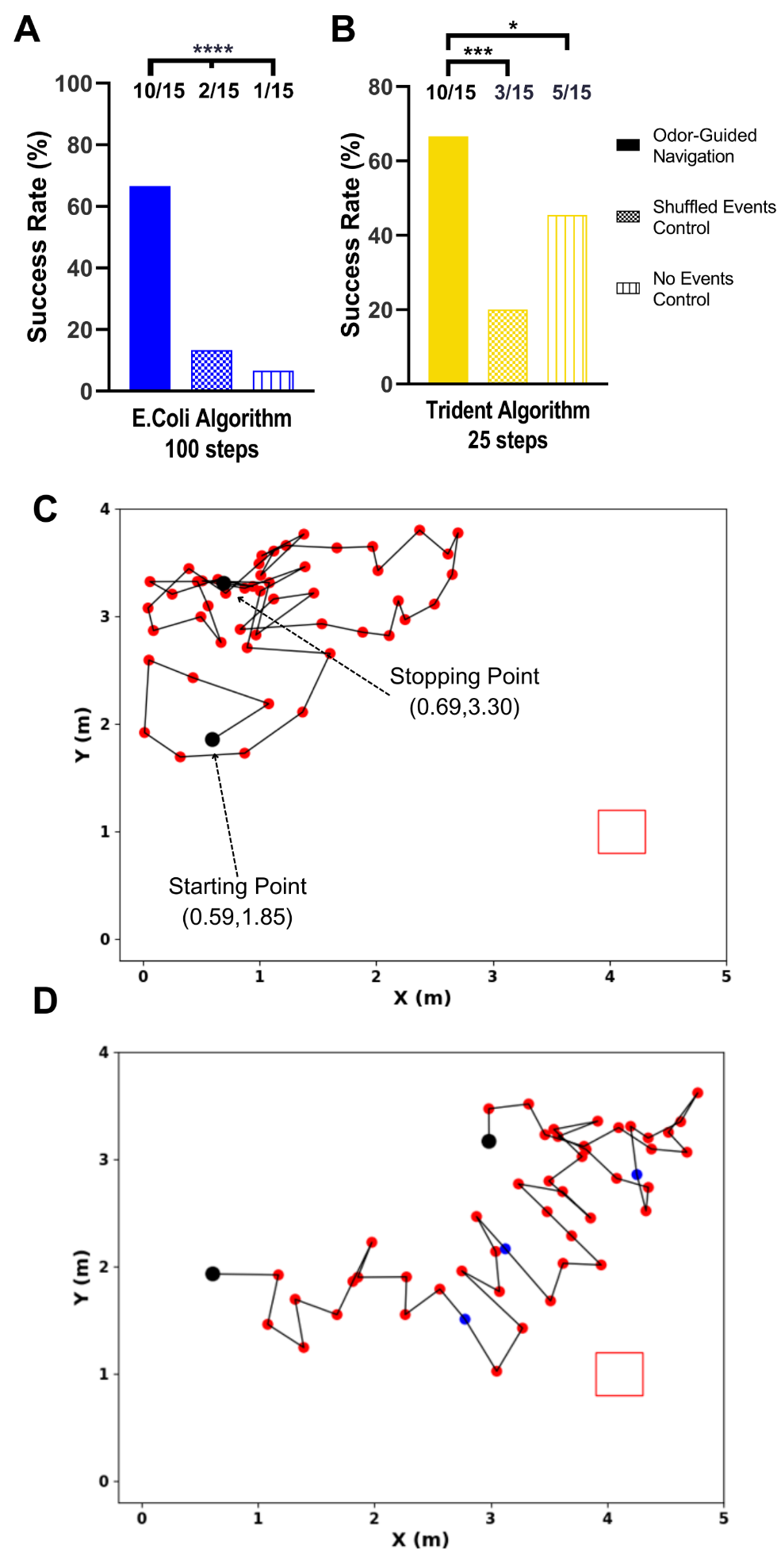
**Suppl. Figure. 5:** **Sniffbot’s Odor Localization Experiments and Controls.** (A) Success rates of odor-guided navigation versus two types of odorless controls for the E. coli algorithm when experiments were limited to 100 steps. Two-tailed binomial test: p<0.0001. (B) Success rates of odor-guided navigation versus two types of odorless controls for the Trident algorithm when experiments were limited to 25 steps. Two-tailed binomial test: p<0.05 (no-event control) and p<0.001 (shuffled event control). (C) Trajectory of the robot in a “no events” control experiment using the E. coli algorithm. (D) Trajectory of the robot in a “shuffled events” control experiment using the E. coli algorithm.

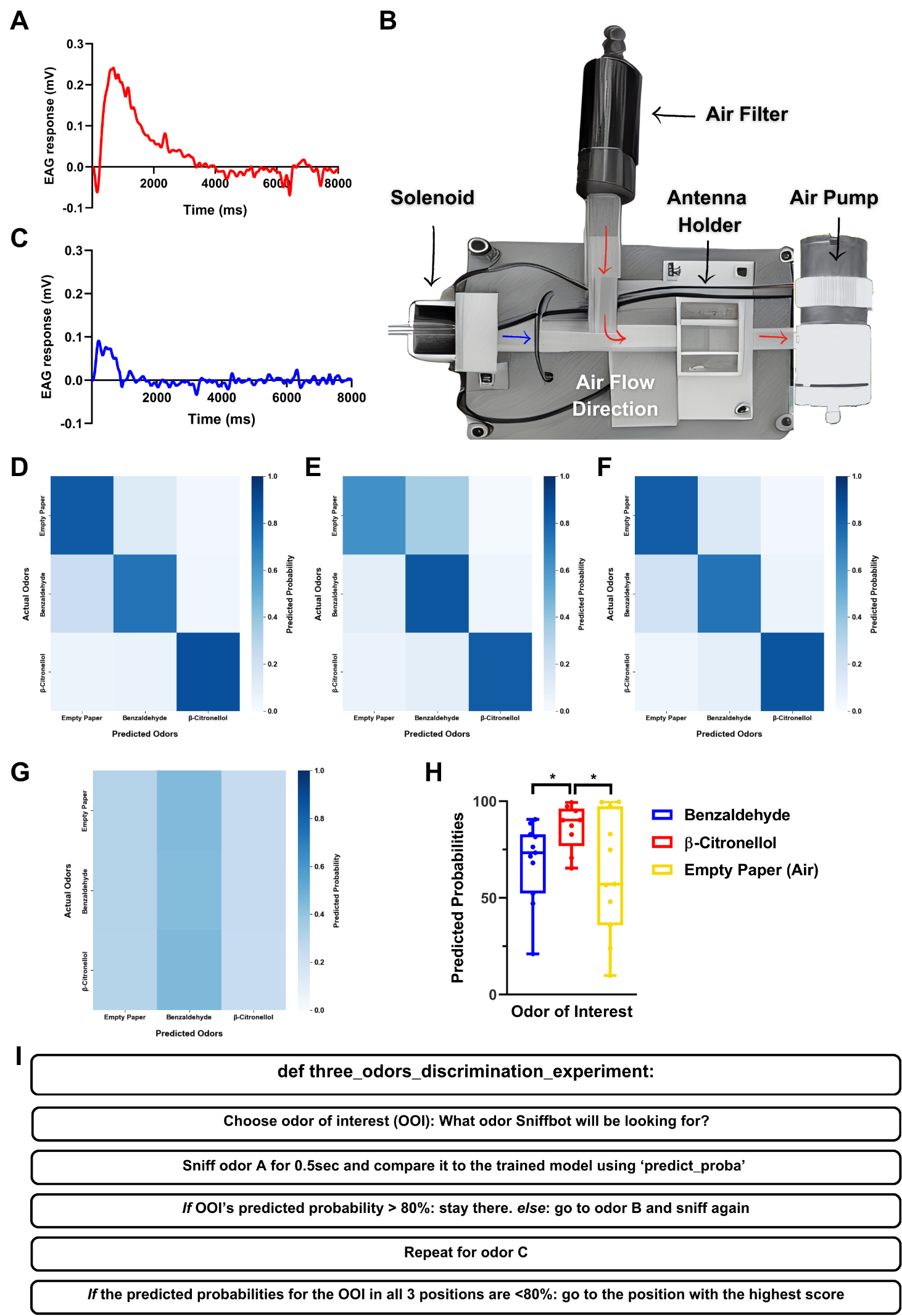

**Suppl. Figure. 6: Sniffbot’s Odor Discrimination Experiment. (A)** Example of the antennal response to the air flow burst immediately after the solenoid opens using the first version of the sniffer in an odorless environment. **(B)** Schematic of the second version of the sniffer. See **Improved version of the sniffer** in the **Supplementary Information**). When the solenoid is closed, only filtered air passes through the antenna (red arrows), allowing the antenna to acclimate to the airflow. When the solenoid opens, unfiltered ambient air merges with the filtered airflow and passes through the antenna (blue arrow). **(C)** Example of the antennal response to the wind burst after the solenoid opens using the second version of the sniffer in an odorless environment. **(D, E, & F)** Confusion matrices for the discrimination training dataset (normalized odorant responses as shown in *Figure 5B*) obtained using the random forest, k-nearest neighbors (KNN), and support vector classifier (SVC), respectively, with average accuracies of 81.76%, 70.04%, and 80.82% over 318 samples. **(G)** Control confusion matrix using randomized (shuffled) data with the logistic regression classifier, showing an average accuracy of 33.02% over 318 samples. **(H)** Predicted probabilities for successful experiments from *Figure 5E****.*** N = 12 antennae for each odor of interest. Two-tailed t-test: p<0.05. **(I)** Schematic of the discrimination experimental procedure.

**Supplementary Movies:**

**Movie. 1:** Sniffbot preparation and experimental procedure.

**Movie. 2:** Sniffbot’s navigation experiment using the Trident algorithm, fast-forwarded ×10.

**Movie. 3:** Example trajectory using the E. coli algorithm: blue points indicate odor detection; red points indicate no detection (left panel); corresponding EAG signal at each point shown on the right panel. Peak detection parameters are displayed on the graph.

**Movie 4:** Same as Movie 3, using the Spiral algorithm.

**Movie 5:** Simulated Sniffbot’s navigation in a virtual odor dispersal map using the E.Coli algorithm (successful trial).

**Movie 6:** Same as Movie 5, using the Spiral algorithm.

**Movie 7:** Same as Movie 5, using the Trident algorithm.

**Movie. 8:** Simulated Sniffbot’s navigation in a virtual odor dispersal map using the E. coli algorithm (failed trial).

**Movie. 9:** Same as Movie 8 using the Spiral algorithm.

**Movie. 10:** Same as Movie 8 using the Trident algorithm.

**Movie. 11:** Sniffbot’s discrimination experiment with corresponding recorded EAG signals.
